# Functional consequences of a p53-MDM2-p21 incoherent feedforward loop

**DOI:** 10.1101/2024.06.25.600070

**Authors:** Jamie A. Dean, Jose Reyes, Simone Bruno, Michael Tsabar, Ashwini Jambhekar, Galit Lahav, Franziska Michor

## Abstract

Genetically identical tumor cells can respond heterogeneously to therapy, with a subpopulation of cells often entering a temporarily arrested treatment-tolerant state before repopulating the tumor. To investigate how heterogeneity in the cell cycle arrest protein p21 arises, we imaged the dynamics of p21 transcription and protein expression along with those of p53, its transcriptional regulator, in single MCF-7 cells using live cell fluorescence microscopy. Using this approach, we found that the rate of p21 transcription depends on the change in p53 rather than its absolute level. Through combined theoretical and experimental modeling, we determined that p21 transcription is governed by an incoherent feedforward loop mediated by MDM2. This network architecture facilitates rapid induction of p21 expression and variability in p21 transcription. Abrogating the feedforward loop overcomes rapid S-phase p21 degradation, with cells transitioning into a quiescent state that transcriptionally resembles a treatment-tolerant persister state. The potential implications of our findings warrant consideration in the design of therapeutic strategies based on activating p53.

## Introduction

Genetically identical populations of cells often exhibit heterogeneous fates in response to the same environmental exposures. Such non-genetic variability in cell fates represents a major challenge for cancer therapy aiming to eliminate all tumor cells, facilitating the survival of some cells in an otherwise treatment-sensitive population (1). This non-genetic resistance can occur through a subpopulation of cells transiently entering a cell cycle-arrested state, during which they are able to tolerate therapies that target rapidly proliferating cells (2–5), including radiation and chemotherapy, before re-entering a proliferative state to repopulate the tumor. So-called cancer “persister” cells are now increasingly thought to be a substantial source of treatment failures in oncology and, hence, there is a pressing need to develop novel treatment approaches to overcome this form of resistance. To realize these new therapeutic strategies, an improved understanding of non-genetic resistance mechanisms is required. Knowledge of mechanisms underlying variability in cell cycle phase transitions could also inform approaches to protect normal cells from anti-cancer therapies and reduce treatment side effects.

Previous work has shown that variability in cell fates can be explained by variability in the dynamics of transcription factors regulating fate-determining genes. Different stimuli can give rise to differences in the dynamics of the same transcription factor and result in divergent cell fates. For example, the levels of p53, a key transcription factor regulating the response to DNA damage, oscillate in response to ionizing radiation (IR) cells, but exhibit a sustained increase in response to ultraviolet (UV) radiation (6). These different patterns of p53 expression dynamics are associated with different fates: senescence and apoptosis, respectively. However, even when cells are exposed to the same treatment and exhibit the same broad patterns of transcription factor dynamics, subtle variability in those dynamics can explain cell fate variability. Such subtle non-genetic intercellular heterogeneity in the dynamics of p53 can explain whether cells die or survive following treatment with cisplatin chemotherapy (7).

Cell cycle arrest is primarily controlled by the cyclin-dependent kinase inhibitor, *CDKN1A* (p21), which is transcriptionally activated by p53 in response to DNA damage (8–12). Entry and exit of cell cycle arrest is controlled by a bistable switch created by double negative feedback between p21 and CDK2 (13, 14). Noise in p53 protein levels propagates to p21 protein levels and plays a role in cell fate heterogeneity, in particular the ability of cells to enter and escape from cell cycle arrest (13, 15, 16).

Given the importance of p21 expression dynamics in cell fate determination and heterogeneity in p21 dynamics on non-genetic variability in treatment response (17), we sought to quantitatively characterize p21 expression dynamics through integrating single cell time-lapse fluorescence microscopy of p21 and p53 and mathematical modeling of the regulation of and noise propagation through this circuit.

## Results

### Transcription of p21 is dependent on the change in p53 rather than its absolute level

To quantitatively characterize p53-dependent p21 expression, we used a recently developed experimental system to study the dynamics of the p53 and p21 proteins (using fluorescent reporters p53-CFP and p21-mCherry) together with the dynamics of p21 transcription (using the MS2 system) in single MCF-7 breast cancer epithelial cells (18). We treated cells with IR (a single dose of either 10 Gy (n = 274), 5 Gy (n = 311) or 2.5 Gy (n = 245)) and imaged them with time-lapse microscopy for approximately 2 days (Supplementary Fig. 1a-f). Consistent with previous studies of p53 dynamics in MCF-7 cells (6, 19–23), and other human tumor (24) and normal cell lines (15), p53 protein levels oscillated, with a period of approximately 5.5 h, in response to IR (Fig. 1a and Supplementary Fig. 1a, d, g, j). The p53 oscillations resulted in bursts of p21 transcription (Fig. 1b and Supplementary Fig. 1b, e, h, k) and p21 protein accumulation (Fig. 1c and Supplementary Fig. 1c, f, i, l). In some cells, p21 protein levels rapidly decreased to low levels for several hours at a time (Supplementary Fig. 1i, l), leading to a bimodal distribution of p21 (Supplementary Fig. 1m), consistent with the previously characterized rapid p21 protein degradation in S-phase (25–29). The number of p53 oscillations per cell tended to increase with increasing dose (Supplementary Fig. 1a, d), leading to more bursts of p21 transcription (Supplementary Fig. 1b, e) and accumulation of p21 protein (Supplementary Fig. 1c, f).

**Figure 1:**
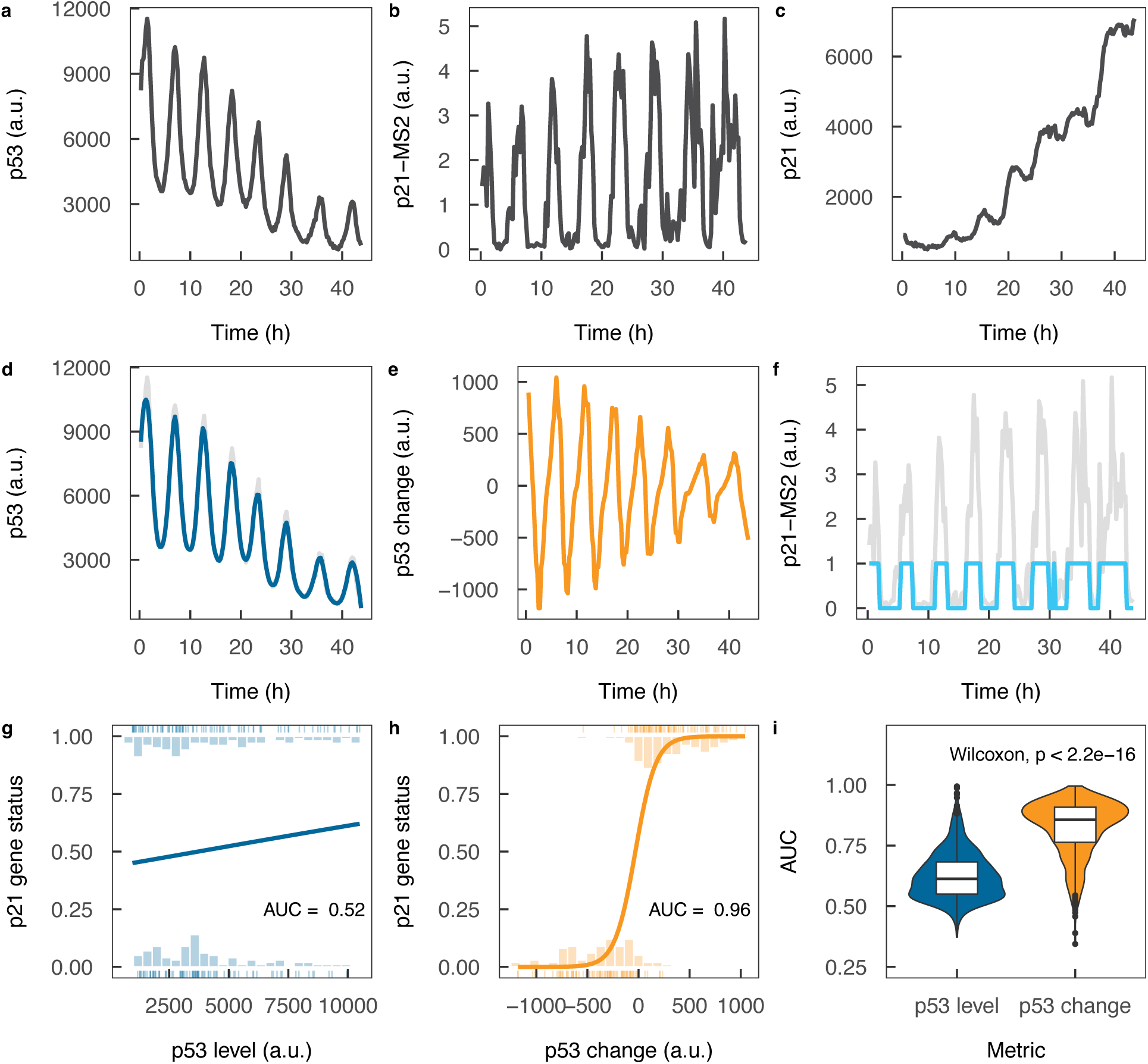
Transcription of p21 is dependent on the change in p53 rather than its absolute level. **(a)** p53 protein dynamics, imaged using cyan fluorescent protein, for a representative cell. **(b)** p21 transcription dynamics, imaged using the MS2 system, for the same representative cell as in (a). **(c)** p21 protein dynamics, imaged using mCherry fluorescent protein, for the same representative cell as in (a). **(d)** Smoothed p53 protein expression (blue line; gray line shows original signal) for the same cell as in (a). **(e)** Change in p53 protein expression for the same cell as in (a). **(f)** Inferred p21 gene state dynamics for the same cell as in (a). The gray line shows the p21-MS2 data from (b) and the pink line shows the fit of the hidden Markov model to the data. The y-axis scale indicates the level of the MS2 signal (gray) and additionally the fit of the two-state hidden Markov model (pink), with a value of 0 indicating the “OFF” state and 1 representing the “ON” state. **(g)** Logistic regression model estimating the binary p21 gene state (in (f)) based on the (detrended) absolute p53 expression level (in (d)). **(h)** Logistic regression model estimating the binary p21 gene state (in (F)) based on the change in the (detrended) p53 expression level (in (e). **(i)** Predictive performance of the logistic regression models based on the absolute p53 expression level and change in p53 expression level for all cells (all dose levels combined; n = 830). AUC = area under receiver operating characteristic curve.

When visualizing the dynamics of p53 and p21 transcription simultaneously in single cells, we observed that p21 transcription occurred only at the times corresponding to the peaks of the p53 oscillations, including instances where p53 oscillation peaks were lower than troughs (Supplementary Fig. 1g, h, j, k). This observation suggested that p21 transcriptional output is dependent on the change in p53 rather than its absolute concentration. To formally test the hypothesis that p21 transcription is dependent on the change in p53, we developed two different statistical models that predict p21 transcription based on p53 protein expression: one based on the absolute level of p53 (Fig. 1d) and a second based on the change in p53 level (Fig. 1e). To infer when the p21 gene was in the “ON” and “OFF” states, we fit a two-state hidden Markov model to the p21-MS2 time course data for each cell, estimating for each time point whether the gene is in the “ON” or “OFF” state (Fig. 1f). We then fit logistic regression models of the inferred p21 gene state based on either the absolute p53 level (Fig. 1g), or the change in p53 level (Fig. 1h). We subsequently compared the predictive performance of the models for each cell, as assessed by the area under the receiver operating characteristic curve (AUC). We found that while the models based on the absolute p53 levels were able to partially predict p21 transcription (mean AUC = 0.66), the models based on the change in p53 were substantially superior (mean AUC = 0.87; (Fig. 1i; p < 2.2E-16, paired Wilcoxon test).

We considered the possibility that long-term trends of increasing and decreasing p53, which are present in the signal, may represent technical artifacts rather than true p53 dynamics and thereby may be confounding our interpretation of p53-dependent p21 transcription. Therefore, to test whether our finding was robust to removal of these long-term trends, we detrended the raw p53 dynamics data (Supplementary Fig. 1n, o) before repeating our analysis. Again, we found that the change in p53 level was a superior predictor of p21 transcription (mean AUC = 0.82) than the absolute p53 level (mean AUC = 0.63; Supplementary Fig. 1p; p < 2.2E-16, paired Wilcoxon test), further supporting our interpretation.

We also repeated this analysis using data from an additional (previously published (18)) experiment employing the same reporter system, but a shorter time course of 15 h and a higher temporal resolution of 2 minutes, to minimize the increasing and decreasing p53 trends apparent in the original experiment. Again, p21 transcription was significantly better predicted by the change in p53 (mean AUC = 0.87) than the absolute p53 level (mean AUC = 0.66; Supplementary Fig. 1q; p < 2.2E-16, paired Wilcoxon test). These findings suggest that p21 transcription is not solely dependent on p53 concentration, but is instead under the control of a more complex regulatory mechanism capable of parsing the change in p53 expression.

### Transcription of p21 is governed by an incoherent feedforward loop that enables p53 change detection

We next investigated how the change in p53 could be parsed by the p21 promoter. Change detection cannot be achieved by a two-node network, such as a network consisting of only a p53 node and a p21 node; instead, the simplest gene network capable of change detection is a three-node network. Theoretical work has shown that very few three-node networks (0.1% of 0.5 million circuit topologies) are capable of conferring change detection (30). Even among these rare networks capable of fold-change detection, only three have been observed experimentally: the incoherent type-1 feedforward loop (IFFL) (31), the nonlinear integral feedback loop, and the logarithmic sensor (32). These observed architectures achieve the optimal performance in two important trade-offs: the balance between response speed and amplitude, and the balance between noise resistance and amplitude (30), potentially explaining their occurrence in nature. Nonlinear integral feedback loops enable fold-change detection in chemotaxis (33, 34), while logarithmic sensors may play a role in allosteric regulation (35). The IFFL architecture is one of the most frequently occcuring network motifs in gene regulatory networks across different organisms (36–41), suggesting recurrent selection of this network motif in the regulation of gene transcription. Therefore, we focused our analysis on determining whether regulation of p21 transcription is governed in this way. We observed that p21 transcription displayed a rapid switching on and off of p21 transcriptional bursting as opposed to a gradual “ramp up” and “ramp down” (Fig. 1b, Supplementary Fig. 1e, g). This behavior suggests that the transcription of p21 is governed by a molecular titration mechanism, whereby p21 is transcribed when the concentration of p53 is above that of a negative regulator of p21 transcription. As IFFLs are capable of molecular titration and have previously been identified in gene regulation, we focused on this motif as a potential alternative mechanism of p21 transcription to simple positive regulation (PR) by p53.

To formally compare the alternative mechanisms of p21 transcription - PR and IFFL, we constructed stochastic mathematical models of each (Fig. 2a, b; Methods). The key difference between the two models is that in the PR model, the p21 transcription rate increases with increasing p53, while in the IFFL model, the p21 transcription rate increases with an increasing difference between the levels of p53 and the transcriptional repressor, and the transcription rate is 0 when the level of transcriptional repressor exceeds the level of p53. For these models we did not commit to an explicit functional form to describe the p53 dynamics to minimize the number of assumptions; instead, we used the p53 data as input into the models. We simulated p21 transcription dynamics from our two stochastic models for each single cell in our data using the Gillespie stochastic simulation algorithm (Fig. 2c, d) and compared the model simulations to our data by computing the p21-MS2 autocorrelation function (Fig. 2e, f) and the p53 - p21-MS2 cross-correlation function (Fig. 2g, h). This approach takes advantage of the fact that our experimental data captures the transcription factor and transcription dynamics over time in the same single cells. We fit both models to our experimental measurements using approximate Bayesian computation-based inference, repeating the stochastic simulations for all cells for 5,000 combinations of different parameter values, to determine which model could best explain the data (Methods). We found that none of the correlation functions generated by the PR model simulations were able to match the correlation functions observed in the data from the long (Fig. 2i, j, Supplementary Fig. 2a, b, e, f) or short time course experiments (Supplementary Fig. 2i, j), whereas the IFFL model recapitulated these correlation functions well (Fig. 2k, l, Supplementary Fig. 2c, d, g, h). Our Bayesian model selection procedure indicated that the IFFL model provided the best fit to the data for all dose levels and datasets (posterior probability of the IFFL model and not the PR model = 1.0, 1.0, 0.90 and 1.0 for the 10 Gy, 5 Gy, 2.5 Gy long time course datasets and short time course dataset, respectively). An extended version of the PR model, incorporating nonlinear activation of p21 transcription by p53 with a Hill function (PR-Hill Function; Methods), was still unable to recapitulate the experimental correlation functions, confirming the robustness of our model selection (Supplementary Fig. 3). Thus, our combined experimental and theoretical modeling approach suggests that, among the models tested, transcription of p21 is not governed by simple PR by p53, but instead by an IFFL involving p53 and an additional transcriptional repressor.

**Figure 2:**
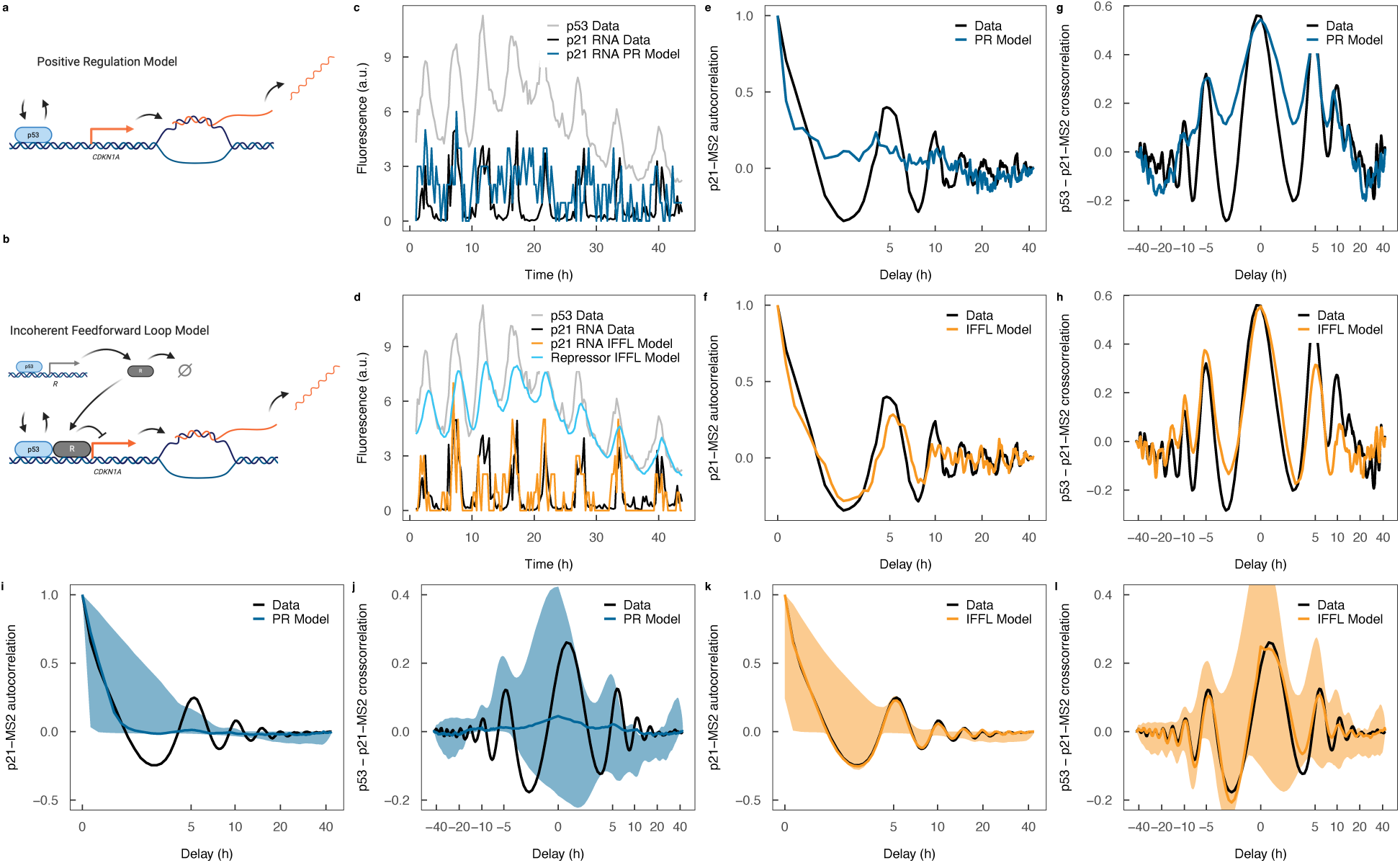
Transcription of p21 is governed by an incoherent feedforward loop that enables p53 change detection. **(a)** Diagram of the positive regulation (PR) model of p21 transcriptional regulation. Switching of the *CDKN1A* gene from the “OFF” to the “ON” state depends solely on p53 bound to the promoter. Created with BioRender.com. **(b)** Diagram of the incoherent feedforward loop (IFFL) model of p21 transcriptional regulation. p53 induces expression of a repressor (R) in addition to switching the *CDKN1A* gene from the “OFF” to the “ON” state. Over time, R accumulates and inhibits *CDKN1A* gene activation by binding (either directly or indirectly) to the DNA. Created with BioRender.com. **(c)** Example simulation of p21 transcription dynamics under the PR model. The gray line indicates the p53 dynamics, the black line indicates the measured p21-MS2 dynamics and the blue line indicates the simulated p21-MS2 dynamics. **(d)** Example simulation of p21 transcription dynamics under the IFFL model. The gray line indicates the p53 dynamics, the black line indicates the measured p21-MS2 dynamics and the blue line indicates the simulated p21-MS2 dynamics. **(e)** p21-MS2 autocorrelation function for the same cell as in (C). The black line shows the data and the dark blue line shows the positive regulation model simulation for an example set of parameter values. **(f)** p21-MS2 autocorrelation function for the same cell as in (d). The black line shows the data and the orange line shows the incoherent feedforward loop model simulation for an example set of parameter values. **(g)** p53-p21-MS2 cross-correlation function for the same cell as in (c). The black line shows the data and the dark blue line shows the positive regulation model simulation for an example set of parameter values. **(h)** p53-p21-MS2 cross-correlation function for the same cell as in (d). The black line shows the data and the orange line shows the incoherent feedforward loop model simulation for an example set of parameter values. **(i)** Mean p21-MS2 autocorrelation function for 10 Gy IR-treated cells for the PR model. **(j)** Mean p53-p21-MS2 cross-correlation function for 10 Gy IR-treated cells for the PR model. **(k)** Mean p21-MS2 autocorrelation function for 10 Gy IR-treated cells for the IFFL model. **(l)** Mean p53-p21-MS2 cross-correlation function for 10 Gy IR-treated cells for the IFFL model. In (i) – (l) the colored lines represent the correlation functions from the simulations that are closest to the correlation functions from the data and the shaded areas represent the 95 percentile confidence intervals of the correlation functions from the simulations. Correlation functions are the means of 274 cells treated with 10 Gy IR.

### The p53-MDM2 interaction is necessary for transcriptional repression of p21

We next attempted to identify the repressor of p21 transcription, representing the third node in the IFFL. We reasoned that the transcriptional repressor should fulfil several criteria: (1) it should be a transcriptional target of p53 (necessary for the IFFL); (2) it should bind to a regulatory region of the *CDKN1A* gene (either directly or indirectly) and (3) it should have dynamics similar to those predicted by our computational model (Fig. 2d). Based on fulfilling these criteria, in addition to the fact that MDM2 has previously been shown to perform a role in transcriptional repression of p53 target genes (42–44), including p21 (45–50), we predicted that MDM2 is the transcriptional repressor. We then leveraged our theoretical model to design and analyze an experiment to test this prediction.

Our computational model predicts that inhibiting the activity of the repressor would result in the p21 transcription rate being dependent on the absolute abundance of p53 rather than the change in p53 concentration. Given that MDM2 is recruited to genes through binding to p53 (44), we reasoned that its predicted transcriptional repressor activity could be inhibited by pharmacologically blocking its binding to p53. Previous work has demonstrated that adding nutlin-3a, which inhibits the binding of MDM2 to p53 (51), to IR substantially alters the dynamics of p53, MDM2 and p21 RNA (as assessed by qPCR and RNA-seq analysis) and protein (as assessed by Western blot and mass spectrometry analysis) levels in MCF-7 cells (52), making it a suitable compound to use. To test our model prediction, we analyzed data from an experiment in which MCF-7 cells were treated with 10 Gy IR alone or in combination with 10 *μ*M nutlin-3a (IR + nutlin-3a), and the dynamics of p53 and p21 expression were imaged by time-lapse microscopy (Supplementary Fig. 4) (18). We then compared the PR and IFFL model fits to the data by simulation and approximate Bayesian computation, as described above. We found that for data of the IR-treated cells, the PR model did not provide a good fit (Fig. 3a, b), whereas the IFFL model did (Fig. 3c, d): the Bayesian model selection procedure strongly favored the IFFL model with a posterior probability of the IFFL model (and not the PR model) of 1.0. However, for the cells treated with IR + nutlin-3a, the PR model described the data similarly well as compared to the IFFL model (Fig. 3e-h) and the posterior probability for the IFFL model was substantially reduced to 0.84. Note that the IFFL model reduces to the PR model when the level of the repressor or its effect are set to 0, which likely explains the preference for this model (although to a lesser extent), even when the PR model can recapitulate the correlation functions in the data. This observation validates our model prediction. The PR-Hill Function model produced the same qualitative behaviors as the PR model (Supplementary Fig. 5), demonstrating that the conclusions are robust to the functional form of the activation of p21 transcription by p53.

**Figure 3:**
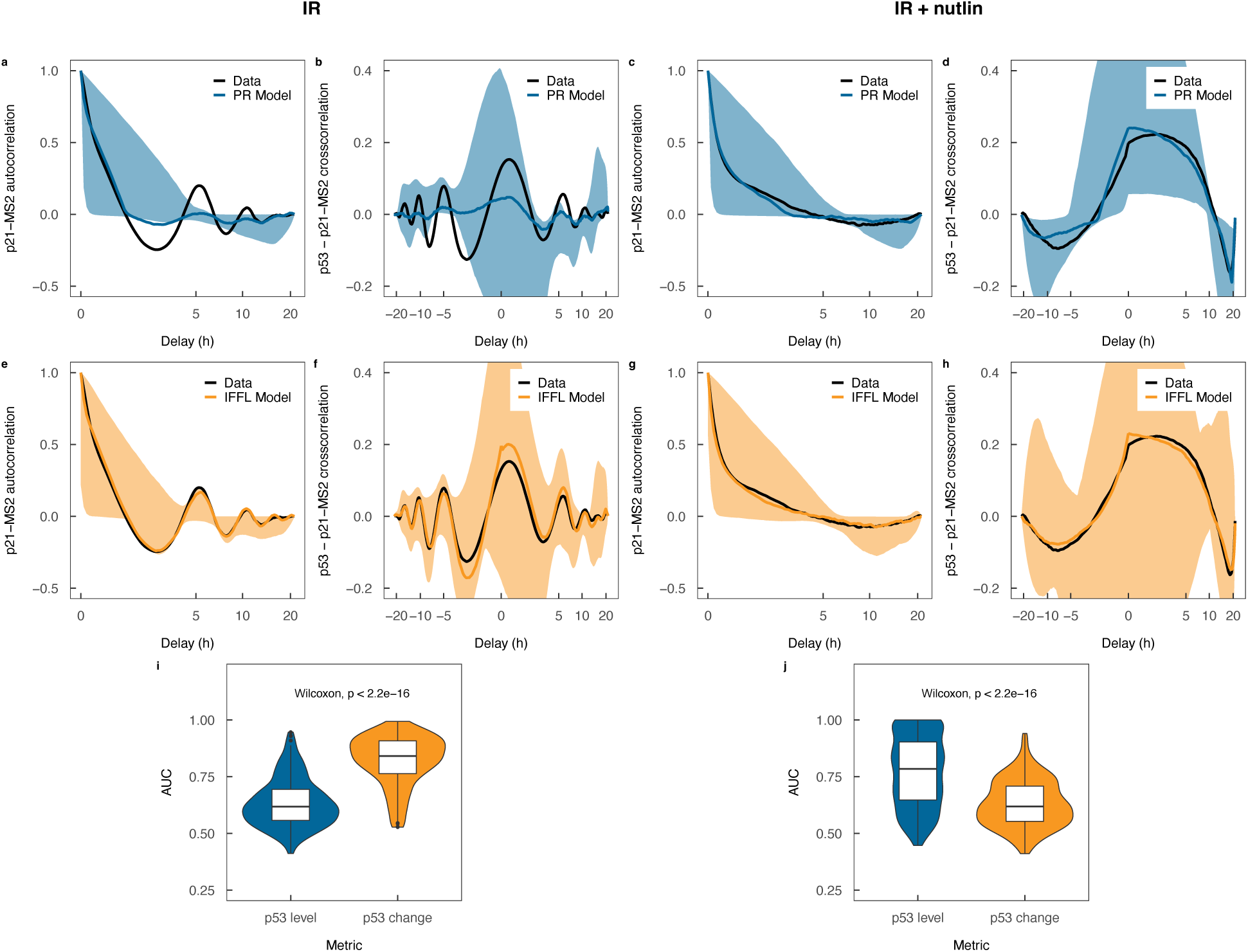
The p53-MDM2 interaction is necessary for transcriptional repression of p21. **(a)** Mean p21-MS2 autocorrelation function for 10 Gy IR-treated cells for the PR model. **(b)** Mean p53 - p21-MS2 cross-correlation function for 10 Gy IR-treated cells for the PR model. **(c)** Mean p21-MS2 autocorrelation function for 10 Gy IR- and 10 *μ*M nutlin-3a-treated cells for the PR model. **(d)** Mean p53 - p21-MS2 cross-correlation function for 10 Gy IR- and 10 *μ*M nutlin-3a-treated cells for the PR model. **(e)** Mean p21-MS2 autocorrelation function for 10 Gy IR-treated cells for the IFFL model. **(f)** Mean p53 - p21-MS2 cross-correlation function for 10 Gy IR-treated cells for the IFFL model. **(g)** Mean p21-MS2 autocorrelation function for 10 Gy IR- and 10 *μ*M nutlin-3a-treated cells for the IFFL model. **(h)** Mean p53 - p21-MS2 cross-correlation function for 10 Gy IR- and 10 *μ*M nutlin-3a-treated cells for the IFFL model. In (a) – (h), the colored lines represent the correlation functions from the simulations that are closest to the correlation functions from the data and the shaded areas represent the 95 percentile confidence intervals of the correlation functions from the simulations. Correlation functions are the means of 248 and 251 cells treated with 10 Gy IR and 10 Gy IR + 10 *μ*M nutlin-3a, respectively. **(i)** Predictive performance of logistic regression models of p21 gene state based on p53 expression level and change in p53 expression level for 10 Gy IR-treated cells (n = 248). **(j)** Predictive performance of logistic regression models of the p21 gene state based on p53 expression level and change in p53 expression level for 10 Gy IR- and 10 *μ*M nutlin-3a-treated cells (n = 251). AUC = area under the receiver operating characteristic curve.

To further interrogate our proposed mechanism of p21 regulation, we returned to our statistical models of p21 transcription. We found that, for the cells treated with IR alone, the model based on the change in p53 was substantially superior (mean AUC = 0.82) to the one based on absolute level of p53 (mean AUC = 0.64), as before (Fig. 3i; p < 2.2E-16, paired Wilcoxon test). However, for the cells treated with IR and nutlin-3a, the reverse was true (Fig. 3j; p53 level: mean AUC = 0.78; p53 change: mean AUC = 0.63; p < 2.2E-16, paired Wilcoxon test). This observation provides further support for our proposed mechanism that the p53-MDM2 interaction is necessary for the transcriptional repression of p21.

### The incoherent feedforward loop may confer beneficial functionality

We then sought to elucidate the functional consequences of p21 transcription being governed by an IFFL. Previous theoretical (53) and experimental (54) studies have proposed that the type-1 IFFL increases the response rate of the transcriptional target gene. The capability to rapidly enact cell cycle arrest upon detection of DNA damage would very likely be advantageous to cells. Therefore, we investigated whether this scenario was the case for p21. Experimentally manipulating a system to test the effects of a specific regulatory interaction in isolation, without affecting other properties of the system such as steady state levels, is extremely challenging. However, such a controlled comparison can be achieved using mathematical modeling (55). We thus employed mathematical modeling to investigate the effects of the IFFL on the p21 response rate. To this end, we modeled the dynamics of p53-dependent p21 protein expression with p21 production governed either by PR or an IFFL (Methods). We selected the p21 production rates for the two models such that the “quasi-steady state” levels of p21, i.e. where the time-averaged p21 level remains constant and does not continue to increase with additional p53 pulses, were approximately equal for both models. Using these models, we found that the time taken for p21 protein expression to reach half the value of its quasi-steady state level for the PR model was approximately 15 h (Fig. 4a) while the corresponding time for the IFFL model was 5 h (Fig. 4b). The higher p21 induction rate for the IFFL model is due to the production rate of p21 being higher than for the PR model during the time taken for the levels of the repressor to accumulate. This observation suggests that the IFFL indeed increases the rate of p21 induction, which would facilitate rapid cell cycle arrest following DNA damage.

**Figure 4:**
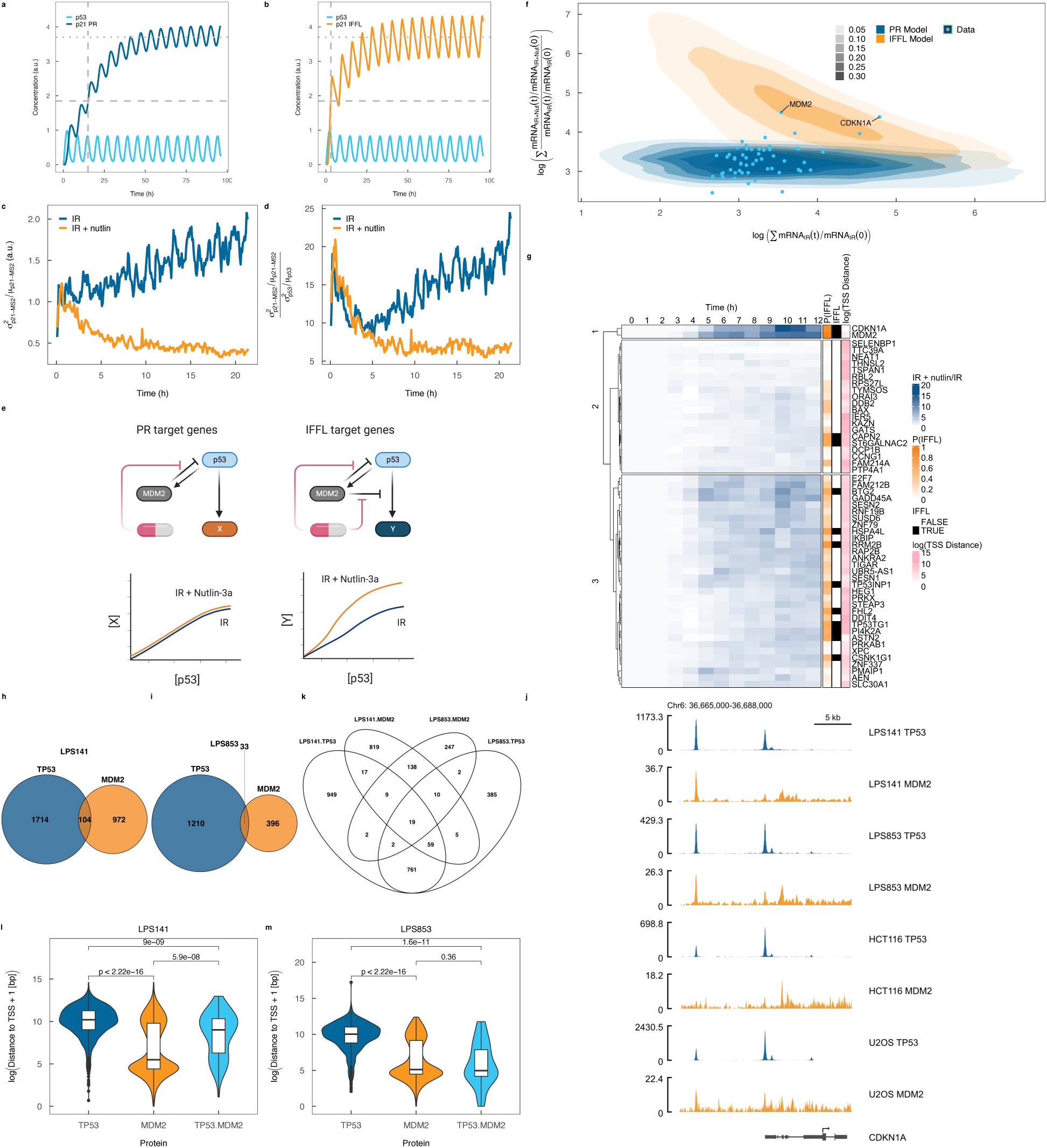
The incoherent feedforward loop increases p21 expression response rate and noise and governs a subset of other p53 target genes. **(a)** Simulated p21 protein dynamics with p21 production governed by PR. **(b)** Simulated p21 protein dynamics with p21 production governed by an IFFL. **(c)** Longitudinal measurements of p21-MS2 noise, as measured by the Fano factor, in cells treated with 10 Gy IR (n = 248) and 10 Gy IR + 10 *μ*M nutlin-3a (n = 251). **(d)** Longitudinal measurements of p21-MS2 to p53 protein noise ratio, with noise measured by the Fano factor, in cells treated with 10 Gy IR (n = 248) and 10 Gy IR + 10 *μ*M nutlin-3a (n = 251). **(e)** Approach to determining whether the transcription of p53 target genes are governed by PR or IFFL. Under PR, the increased expression of a gene upon adding nutlin-3a to IR would be due to the increase in p53 levels alone, while under IFFL, the increased expression would be greater than that expected from the increase in p53 levels alone. Created with BioRender.com. **(f)** Positive regulation (dark blue) and incoherent feedforward loop (orange) model simulations for approximate Bayesian computation. The plot shows the relationship between gene expression following 10 Gy IR + nutlin-3a and 10 Gy IR alone for each model and the corresponding measurements for p53 transcriptional target genes (light blue). Measurements are from MCF-7 cells treated with 10 Gy IR or 10 Gy IR + nutlin-3a (59). **(g)** Time course of the ratio of mRNA expression of TP53 transcriptional target genes following 10 Gy IR + nutlin-3a treatment to mRNA expression following 10 Gy IR treatment (blue). For each gene the model-predicted probability that its expression is governed by an IFFL (P(IFFL); orange), classification of whether P(IFFL) is greater than 0.5 (black), and the distance from the TP53 binding site to the transcription start site (TSS Distance; pink) are also indicated. Data are from the same experiment as in (f). **(h)** Venn diagram of TP53 and MDM2 ChIP-seq peaks in the LPS141 liposarcoma patient-derived cell line. **(i)** Venn diagram of TP53 and MDM2 ChIP-seq peaks in the LPS853 liposarcoma patient-derived cell line. **(j)** TP53 and MDM2 ChIP-seq peaks at the *CDKN1A* gene in LPS141, LPS853, HCT116 and U2OS cells. **(k)** Venn diagram of TP53 and MDM2 ChIP-seq peaks in the LPS141 and LPS853 cell lines. **(l)** Distance between TP53, MDM2 and overlapping TP53 and MDM2 (TP53.MDM2) ChIP-seq peaks and transcription start site in the LPS141 liposarcoma patient-derived cell line. **(m)** Distance between TP53, MDM2 and overlapping TP53 and MDM2 (TP53.MDM2) ChIP-seq peaks and transcription start sites in LPS853 liposarcoma patient-derived cell line. P(IFFL) – posterior probability of IFFL model; IFFL - posterior probability of IFFL model greater than 0.5; TSS Distance – distance between the transcription start site and the closest p53 ChIP-seq peak.

It has also been proposed that the type-1 IFFL generates a large extent of noise (56), which would facilitate cells to enact “bet hedging”, a strategy to minimize the risk of population extinction. Consistent with that pattern, p21 has been observed to exhibit substantial heterogeneity in unstressed conditions, reminiscent of bet hedging (14). Comparing the p21-MS2 noise between cells treated with IR and cells treated with IR and nutlin-3a revealed that inhibiting the p53-MDM2 interaction significantly decreased noise in p21 transcription, when noise was measured by either the Fano factor (standard deviation squared/mean) (Fig. 4c) or the coefficient of variation (standard deviation/mean) (Supplementary Fig. 6a). This finding suggests that the IFFL increases p21 noise. However, given that nutlin-3a increases p53 protein stability and reduces p53 protein noise (Supplementary Fig. 6b), the reduction in p21-MS2 noise upon addition of nutlin-3a to IR may be solely due to its effects on p53 noise and not the IFFL. We, therefore, compared the ratio of the p21-MS2 noise to the p53 noise and found that the p21-MS2 to p53 protein noise ratio was higher in cells treated with IR than when the IFFL is abrogated through treatment with nutlin-3a when measuring noise using either the Fano factor (Fig. 4d) or coefficient of variation (Supplementary Fig. 6c). Both the p21-MS2 noise and p21-MS2 to p53 protein noise ratio were consistent between IR-treated cells in the experiments (Supplementary Fig. 6d, e). This observation suggests that MDM2-mediated transcriptional repression of p21 enhances p21 transcriptional noise.

Another potential property of IFFLs is that they enable the transmission of multiple signals in the dynamics of a single transcription factor, known as “signal multiplexing” (57, 58), when integrated into a larger network. The dynamics of a single transcription factor could simultaneously encode two signals: one encoded in its absolute level and the other in the change in its level. Target genes whose transcription is governed by PR would decode the signal encoded in the absolute level and those whose transcription is governed by an IFFL would decode the change in transcription factor level. If multiple signals were indeed encoded in p53 dynamics, then we would expect that to decode these signals, some p53 transcriptional target genes would be regulated by PR and others by IFFL. To test this hypothesis, we performed a new, mathematical model-based analysis of previously published RNA-seq time course data of p53 transcriptional target genes in MCF-7 cells treated with 10 Gy IR or 10 Gy IR and nutlin-3a (59). Our modelling approach was based on distinguishing between differences in mRNA expression of p53 target genes following treatment with IR or IR and nutlin-3a that were due to differences in p53 levels alone and increases in expression greater than what would be expected based on increased p53 level alone (Fig. 4e). We modelled mRNA expression dynamics using ordinary differential equations with transcription governed by either PR or IFFL and performed model comparison individually for each p53 transcriptional target, previously defined based on RNA-seq and TP53 chromatin immunoprecipitation (ChIP)-seq data from MCF-7 cells treated with 10 Gy IR or 10 Gy IR and nutlin-3a (59), using approximate Bayesian computation (Fig. 4f, g; Methods). This analysis indicated a high posterior probability that p21 transcription was governed by a MDM2-mediated IFFL (0.88). The agreement of this finding, made from bulk RNA-seq data, with that made from our single cell time-lapse microscopy data validates the suitability of this approach for determining whether the transcription of genes is governed by a MDM2-mediated IFFL. We found that for 22% of p53 transcriptional target genes, mRNA expression dynamics were best described by a MDM2-mediated IFFL model (posterior probability of IFFL model > 0.5), whereas the remaining 78% were best described by a PR model (Fig. 4f, g). In addition to p21, the IFFL model was strongly preferred for MDM2 (posterior probability = 0.91). This analysis provides evidence that, for a subset of genes, the addition of nutlin-3a to IR results in gene expression changes that are not solely dependent on changes in p53 dynamics, but also from the alleviation of the MDM2-dependent transcriptional repression of p53 target genes. These findings are consistent with a gene regulatory network structure that can decode a multiplexed signal encoded in p53 dynamics.

To determine whether similar MDM2-mediated transcriptional repression may occur in a subset of p53 target genes in other cell lines, we reanalyzed TP53 and MDM2 ChIP-seq data of four different (untreated) cell lines from a recent study (60), searching for overlapping TP53 and MDM2 peaks. Such overlapping peaks are likely necessary for MDM2-mediated repression of p53-dependent transcription. Consistent with IFFL regulation of a subset of p53 transcriptional target genes, we found both overlapping and non-overlapping TP53 and MDM2 peaks in all of the cell lines (Fig. 4h, i and Supplementary Fig. 6f, g). Both the number of overlapping TP53 and MDM2 peaks and the total number of MDM2 peaks were higher in LPS141 (104 overlapping peaks, 1076 MDM2 peaks) and LPS853 cells (33 overlapping peaks, 429 MDM2 peaks) than HCT116 (3 overlapping peaks, 142 MDM2 peaks) and U2OS cells (4 overlapping peaks, 86 MDM2 peaks), likely due to the fact that these cell lines harbor MDM2 amplifications. LPS141 has higher MDM2 mRNA expression (2043 TPM) than LPS853 (802 TPM) (60), which could explain the increased number of MDM2 peaks and TP53-MDM2 overlapping peaks in LPS141 cells. Three of the 4 cell lines (LPS141, LPS853, U2OS) had overlapping TP53 and MDM2 peaks on the *MDM2* (Supplementary Fig. 6h) gene and 2/4 (LPS141, LPS853) had overlapping TP53 and MDM2 peaks on the *CDKN1A* gene (Fig. 4j). These observations suggest that the expression of a subset of p53 target genes may be regulated by an MDM2-mediated IFFL, with the IFFL-regulated genes differing between different cells.

To determine whether factors beyond MDM2 expression could determine cell line-specific TP53-MDM2 cobinding, we compared data of the two MDM2 amplified cell lines (LPS141, LPS853), which had sufficient TP53-MDM2 cobinding sites to allow for statistical comparisons. When investigating the overlap of TP53-MDM2 cobinding sites between cell lines, we found that 42% (14/33) of cobinding sites in LPS853 cells, which have lower MDM2 expression, were not present in LPS141 cells (Fig. 4k). Therefore, differences in TP53-MDM2 cobinding between cell lines cannot be explained by differences in MDM2 expression level alone. Of the 14 TP53-MDM2 cobinding sites unique to LPS853 cells, 2 (14%) were bound by TP53 alone in LPS141 cells, suggesting that cell line-specific differences in TP53-MDM2 cobinding were predominantly driven by differential TP53 binding. To understand whether differences in histone modifications could explain the differences in TP53-MDM2 cobinding sites between the cell lines, we analyzed H3K27ac ChIP-seq data. In both cell lines we found that TP53-MDM2 cobinding sites overlapped with H3K27ac peaks substantially more frequently (LPS141: 100/113 = 88%; LPS853: 35/37 = 95%) than sites bound by TP53 alone (LPS141: 456/1701 = 27%; LPS853: 470/1197 = 39%) (Supplementary Fig. 6i). Of the 14 LPS853 TP53-MDM2 cobinding sites not bound by TP53 in LPS141 cells, 11 (79%) overlapped H3K27ac peaks, indicating that most have the potential to be bound by transcription factors. Additionally, 10/14 sites (71%) were bound by MDM2 alone, suggesting that MDM2 may be recruited to chromatin by other cofactors beyond TP53, as previously observed (61, 62), the expression of which could vary between cell lines. Taken together, these findings suggest that cell type-specific TP53-MDM2 co-binding of p53 target genes may be determined by a combination of MDM2 expression level, histone modifications and co-factor expression.

When analyzing the MCF-7 cell TP53 ChIP-seq data, we aimed to understand the factors determining which p53 target genes are regulated by an IFFL. We noted that genes inferred to be IFFL-regulated with high probability, *CDKN1A* and *MDM2*, had a short distance between the TP53 binding site and transcription start site (Fig. 4g). We searched for a similar relationship in the LPS141 and LPS853 TP53 and MDM2 ChIP-seq data and found that the TP53 binding sites for genes bound by both TP53 and MDM2 were significantly closer to the transcription start sites (predominantly in regions designated promoters) than for genes bound by TP53 alone (predominantly in regions designated enhancers) (LPS141 p = 9.0E-9, LPS853 p = 1.6E-11; Fig. 4l, m). LPS141 cells had a higher fraction of TP53-MDM2 cobinding sites localized further from transcription start sites than LPS853, which could potentially be due to increased MDM2 expression (and MDM2 ChIP-seq peaks) in this cell line. Co-binding of TP53 and MDM2, facilitating transcriptional repression, may therefore preferentially occur at gene promoters rather than enhancers. This finding suggests that while the set of genes cobound by TP53 and MDM2 varies between cell lines, there may be an increased propensity for a subset of p53 target genes to be transcriptionally repressed by MDM2 based on the locations of their p53 binding sites.

### Abrogating MDM2-mediated transcriptional repression of p21 prevents the G1-S transition

Therapeutically targeting the interaction between p53 and MDM2 is an active area of research in oncology. This strategy is based on increasing p53 levels of p53 wild-type cancer cells through inhibiting its MDM2-mediated degradation. However, the distinct effects of abrogating the p21 IFFL by inhibiting the p53-MDM2 interaction have not been studied. We therefore investigated the impact of nutlin-3a on p21 expression in more detail. To disentangle the effects of inhibiting MDM2-mediated p53 degradation and MDM2-mediated p21 transcriptional repression on the accumulation of p21, we developed a combined theoretical and experimental modeling approach.

Theoretical modelling of p21 expression dynamics (Methods) predicts that at quasi-steady state with high concentrations of p53, the concentration of p21 is given by 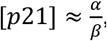 where *α* and *β* are the production and degradation rates of p21, respectively. This relationship indicates that if nutlin-3a only increases p53 levels, without affecting the p21 production rate per unit p53, then when [p21] is plotted against [p53] the y-intercepts of IR- and IR + nutlin-3a-treated cells will be the same. Alternatively, if nutlin-3a increases the production rate of p21 per unit p53, then the y-intercept of IR + nutlin-3a-treated cells will be greater than IR-treated cells. Beyond detecting qualitative differences, the equation enables the ratio of p21 production and degradation rates to be inferred from measurements of p21 concentration. To test the validity of this theory, we investigated experimentally derived values of [p21] and [p53] using the data from cells treated with IR and IR + nutlin-3a (Supplementary Fig. 4). We used the data acquired from the final 2.5 h of the experiment (∼19 h – 21.5 h following IR), to allow sufficient time for p21 levels to reach a quasi-steady state, and restricted our analysis to cells for which the p21 quasi-steady state had been achieved (Supplementary Fig. 7a-f). The data clustered into two populations of cells for both conditions (Fig. 5a). This bimodal distribution is expected as p21 has two different degradation rates, with a rapid degradation occurring during S-phase and a slower degradation during the remainder of the cell cycle, leading to different steady state levels. Indeed, as predicted by the theory, this analysis gave rise to straight lines with slopes of 0 for both conditions (Fig. 5a, b). The y-intercepts of the lines differed, with the lines corresponding to the cells treated with IR + nutlin-3a having greater y-intercepts (Fig. 5a, b). However, there is large uncertainty in the estimate of the y-intercept (and slope) of mixture component 1 (lower steady state level of p21) for cells treated with IR + nutlin-3a, which may be at least partially explained by the small number of cells contributing to this estimate. This observation suggests that the addition of nutlin-3a increases the production rate of p21 substantially beyond what would be expected based on the increased levels of p53 alone. This finding provides further evidence to support our finding that MDM2 represses p53-dependent p21 transcription.

**Figure 5:**
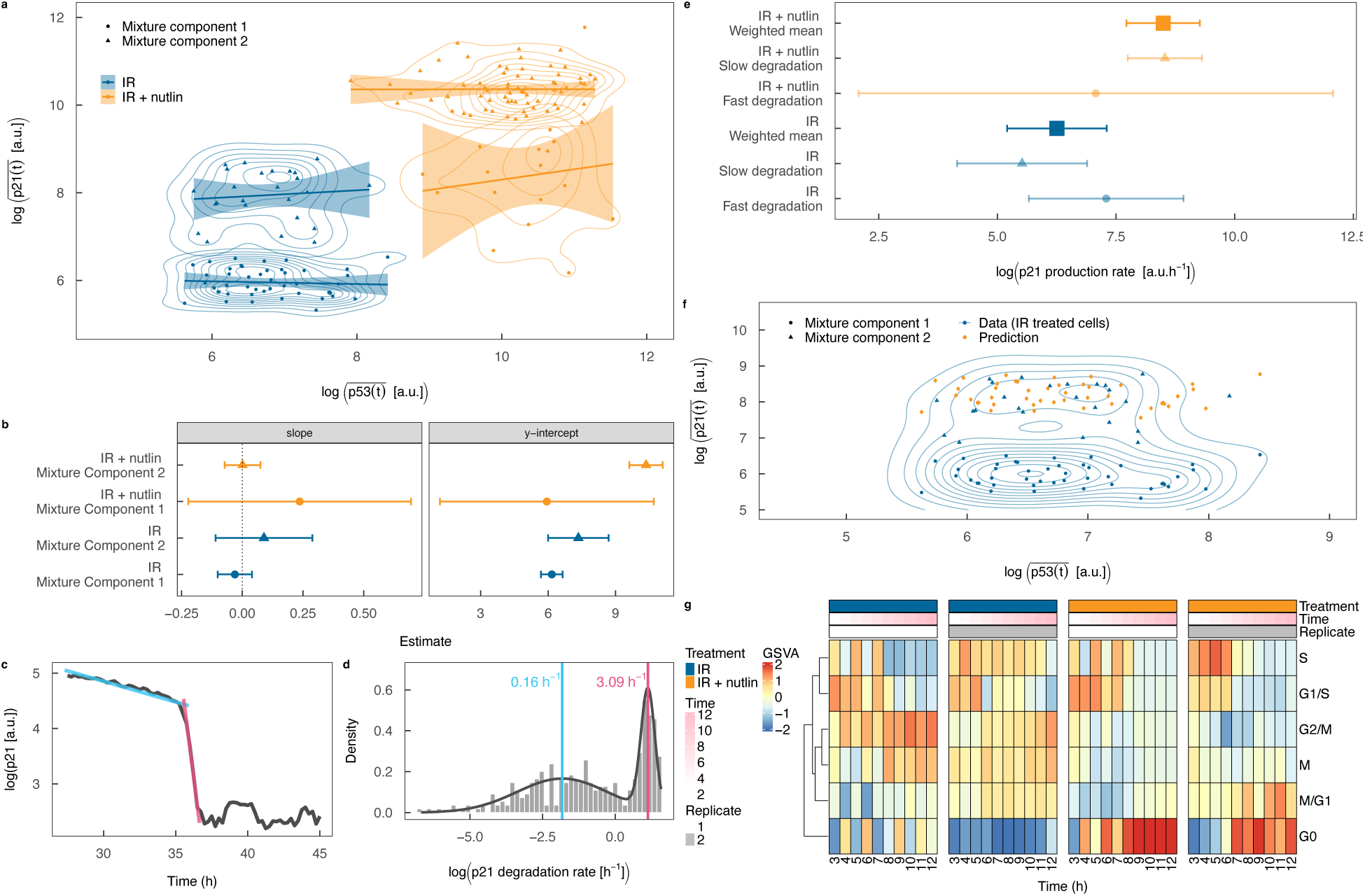
Abrogating MDM2-mediated transcriptional repression of p21 prevents the G1-S transition. **(a)** Relationship between p21 protein and p53 protein averaged over time points at quasi-steady state for single cells treated with 10 Gy IR (n = 248) or 10 Gy IR + 10 *μ*M nutlin-3a (n = 251). The circular and triangular points designate the first and second components of the mixture distributions, respectively. The contours show the distributions of the data. The lines show linear regression fits to the data. **(b)** Slopes and y-intercepts of the regression lines in (a). The dotted line indicates the value of the slope predicted by the mathematical model. **(c)** Fit of the piecewise linear regression model to p21 protein dynamics for a representative example cell following addition of 40 nM p21 siRNA to 10 Gy IR-treated cells to infer p21 protein degradation rates. **(d)** Distribution of inferred p21 protein degradation rates for all cells (n = 345). The histogram shows the inferred degradation rates, the gray line shows a fit of a Gaussian mixture model to the rate distribution and the light blue and pink vertical lines indicate the means of the slow and fast degradation rates across all cells. **(e)** Inferred p21 protein production rates for MDM2-repressed and unrepressed p21 production, inferred from the inverse variance weighted mean of the inferred production rates in cells with high and low p21 degradation rates. **(f)** Prediction of p21 protein levels from abrogating the IFFL without altering p53 protein levels in IR-treated cells in S-phase. The orange points (and error bars) show the predictions (and their standard errors) and the blue points (and contours) show the data (and distribution) for IR-treated cells (same as in (a)). Abrogating the IFFL without altering p53 levels is predicted to increase p21 levels of S-phase cells to the levels of those in non-S-phase cells. **(G)** Gene set variation analysis of cell cycle phase transcriptional signatures (65) applied to RNA-seq data measured hourly from 3 – 12 h following treatment of MCF-7 cells with 10 Gy IR or 10 Gy IR + nutlin-3a. IR – ionizing radiation; GSVA – gene set variation analysis.

To this point our theory allowed us to infer the ratio of the production and degradation rates of p21. Quantifying the p21 production rate, in the presence and absence of nutlin-3a, enables further insights to be gleaned. In particular, previous theoretical models have suggested that the G1-S transition can be prevented by the accumulation of p21 in S-phase, achieved through loss of Cdt2 (63, 64), part of the CRL4^Cdt2^ ubiquitin ligase complex, which is involved in rapid degradation of p21 during S-phase (25–28). We considered that p21 accumulation during S-phase, and hence prevention of the G1-S transition, could also be achieved through increasing its production rate to a level higher than its degradation rate. Therefore, we performed an additional experiment to measure the p21 degradation rate, which would enable inferring the p21 production rates in the context of treatment with IR and IR + nutlin-3a. To measure the rates of p21 protein degradation in single cells, we exposed cells to siRNA targeting either p21 alone or p53 and p21 at 20 h following IR (Supplementary Fig. 7g). This perturbation led to a two-phase decay of p21 in single cells (Fig. 5c). Initially p21 levels decreased slowly, followed by a rapid decrease, presumably due to cells entering S-phase during which CRL4^Cdt2^ ubiquitin ligase complex-dependent rapid degradation occurs (25–27). We inferred these two different decay rates for all cells individually (from both conditions) by fitting a piecewise linear regression model to the log-transformed expression level data for that cell (Fig. 5c). The distribution of p21 degradation rates over all cells was bimodal with peak degradation rates (inferred by Gaussian mixture modeling) of 0.16 h^-1^ and 3.09 h^-1^ (Fig. 5d), corresponding to half-lives of 4.3 h and 0.2 h. These degradation rates imply, based on the equation for steady state levels of p21, that the production rates under IR and IR + nutlin-3a treatments are 520 a.u.h^-1^ and 4880 a.u.h^-1^, respectively (Fig. 5e).

Having inferred the two different p21 production and degradation rates allowed us to mathematically decouple the effects of MDM2 on p53 levels and p21 production per unit concentration of p53 by computing the levels of p21 under different combinations of production and degradation rates. This analysis suggests that inhibiting MDM2-mediated p53 degradation alone would be insufficient to cause p21 accumulation during S-phase (Supplementary Fig. 7h). However, additionally inhibiting the transcriptional repression of p21 achieves p21 accumulation in S-phase (Fig. 5f). This finding implies that the pharmacological abrogation of the p21 IFFL leads to cells remaining in or transitioning to the G1- or G0-phases of the cell cycle and that this would not be achieved through increasing p53 alone without abrogating the IFFL.

To test this prediction, we computed cell cycle signature (65) scores for previously published RNA-seq time course data of cells treated with IR alone or IR + nutlin-3a (59) using gene set variation analysis. This analysis indicated that cells treated with IR predominantly arrest at the G2/M checkpoint whereas cells treated with IR + nutlin-3a exit the cell cycle, arresting in G0 (Fig. 5g). We reasoned that this dramatic effect on cell cycle progression should also lead to changes in cell morphology associated with cell cycle phase, such as nuclear area. In cells that were treated with IR, the nuclear area increased over time following treatment (p < 2E-16, paired t-test; Supplementary Fig. 7i, j), suggestive of cell cycle progression. However, in cells that were treated with IR + nutlin-3a, the nuclear area decreased (p = 7.2E-6, paired t-test; Supplementary Fig. 7i, j). This observation provides morphological evidence, in addition to the transcriptional evidence, of the divergent fates of cells treated with IR and IR + nutlin-3a. We corroborated this morphological change by confirming that nuclear size transcriptional signatures were decreased following treatment with IR + nutlin-3a, but not IR (Supplementary Fig. 7k). Together these findings suggest that IR causes cells to undergo G2/M arrest, but abrogating the p53-MDM2 interaction alters the fate of the cells to instead enter a G0 arrest. While the effects of MDM2 on p53 degradation and transcriptional repression are challenging to experimentally decouple, our theoretical modeling supports the conclusion that the divergence in cell cycle progression is specifically related to MDM2-dependent transcriptional repression, rather than increased levels of p53 alone.

### Abrogating the p53-MDM2 interaction steers cells into a persister state

Having shown that abrogating the IFFL substantially altered p21 dynamics and cell cycle progression, we next sought to investigate the relative contribution of these effects to the global context of p53-mediated changes. We, therefore, compared transcriptome time courses in cells treated with IR in the presence or absence of nutlin-3a using RNA-seq data. Differential expression analyses of mRNA time series data (59) identified 1473 upregulated and 1188 downregulated genes upon addition of nutlin-3a to IR (Fig. 6a). This observation is consistent with our previous work showing p53-dependent increases and decreases in expression of different sets of genes following IR treatment in the same cell line (59). Analysis of the transcriptional regulators of the differentially expressed genes (Methods) indicated that the upregulated genes were enriched for transcriptional targets of p53 and p63 (Fig. 6b), as expected. The downregulated genes were enriched for transcriptional targets of E2F4 and LIN9 (Fig. 6c). E2F4 and LIN9 are members of the DREAM complex that represses transcription of cell cycle genes (66) and whose formation is dependent on p21 (67). This finding suggests that inhibiting the interaction between p53 and MDM2 with nutlin-3a increases the levels of p21, activating the DREAM complex and repressing its target genes. The fact that many of the differentially expressed genes are transcriptional targets of E2F4 and LIN9 suggests that p21 may play a major role in the global gene expression changes resulting from inhibiting the p53-MDM2 interaction.

**Figure 6:**
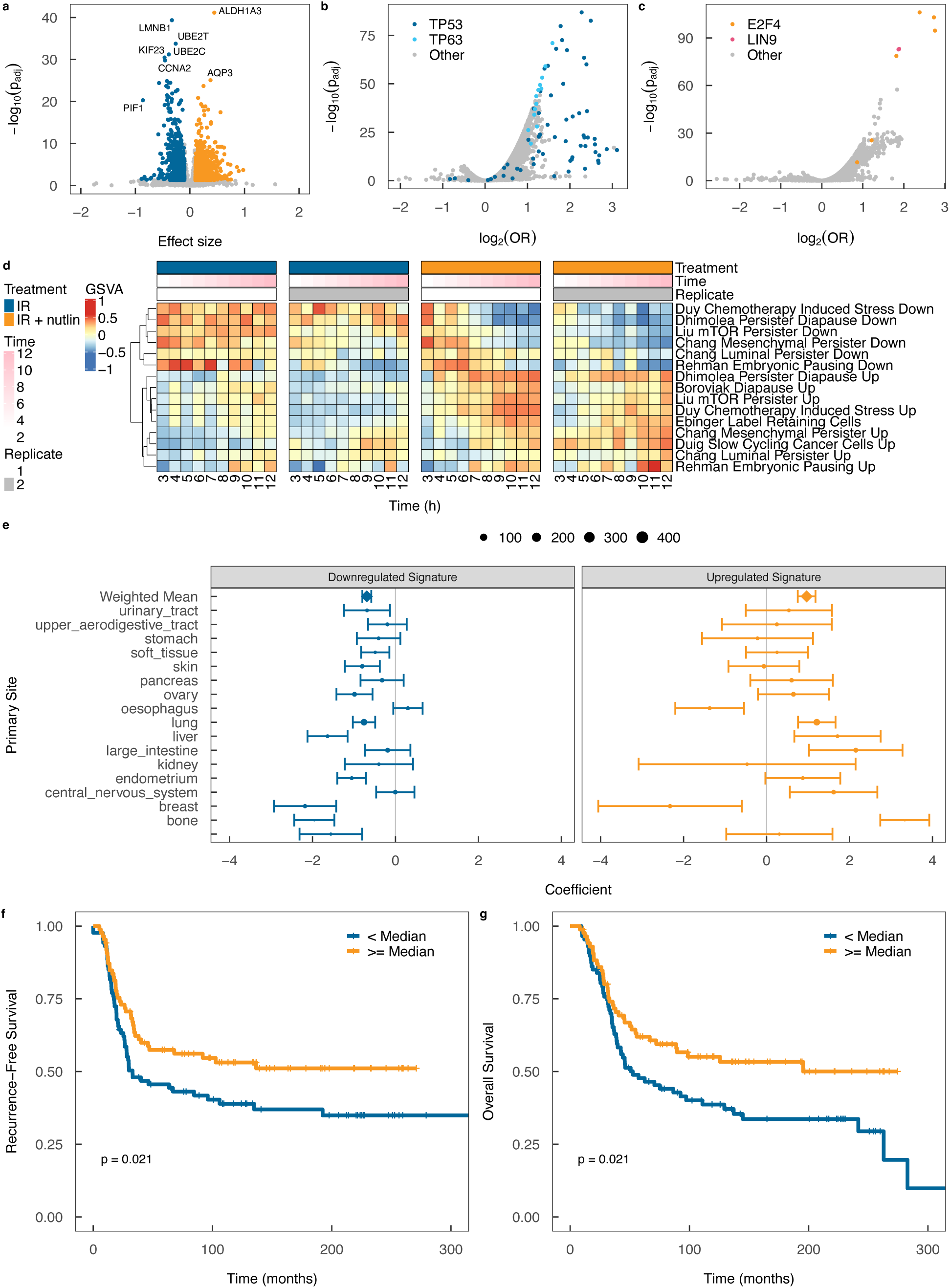
Abrogating the p53-MDM2 interaction steers cells into a persister state. **(a)** RNA-seq differential expression of 10 Gy IR + nutlin-3a- versus 10Gy IR-treated MCF-7 cells with RNA-seq performed hourly from 3 – 12 h following IR. **(b)** Transcriptional regulator enrichment analysis for differentially upregulated genes in 10 Gy IR + nutlin-3a- versus 10 Gy IR-treated MCF-7 cells. **(c)** Transcriptional regulator enrichment analysis for differentially downregulated genes in 10 Gy IR + nutlin-3a- versus 10 Gy IR-treated MCF-7 cells. **(d)** Gene set variation analysis of IR- and IR + nutlin-3a-treated MCF-7 cells with cancer persister cell gene sets^55–62^. **(e)** Linear regression model coefficient evaluating the association between IR response (area under radiation dose-response curve) and GSVA score of genes upregulated and downregulated in 10 Gy IR + nutlin-3a- versus 10 Gy IR-treated MCF-7 cells, in cell lines from the Cancer Cell Line Encyclopedia (71). The weighted mean is the inverse variance weighted mean of the coefficients across all primary tumor sites. The size of the points corresponds to the number of cell lines. **(f)** Association between recurrence-free survival and GSVA score of genes downregulated in 10 Gy IR + nutlin-3a- versus 10 Gy IR-treated MCF-7 cells, in breast cancer patients treated with chemo-radiation therapy from the METABRIC cohort (72) (n = 173). **(g)** Association between overall survival and GSVA score of genes downregulated in 10 Gy IR + nutlin-3a versus 10 Gy IR-treated MCF-7 cells, in breast cancer patients treated with chemo-radiation therapy from the METABRIC cohort (72) (n = 173). The p-values in (F) and (G) are from Cox proportional hazards regression models with the signature score, as a continuous variable, as the covariate. OR – odds ratio; GSVA – gene set variation analysis.

Gene set variation analysis using the Hallmark pathways indicated nutlin-3a-dependent downregulation of E2F and MYC targets and the G2M checkpoint, in addition to the expected upregulation of TP53 (Supplementary Fig. 8a). This observation is consistent with the previously identified antagonistic role of p53 on MYC in breast cancer (68). MYC is also involved in nuclear size regulation, which may explain our earlier finding of nuclear shrinkage upon treatment with IR + nutlin-3a. Downregulation of DREAM complex target genes and MYC is associated with quiescent states such as embryonic diapause, resistance of normal tissues (69) and cancer cells (70) to therapy and treatment-tolerant persister cells. We, therefore, studied persister cell signatures in our RNA-seq data. We found that, for almost all persister cell signatures, IR + nutlin-3a increased upregulated persister cell signatures and decreased downregulated persister cell signatures (Fig. 6d). This finding suggests that adding nutlin-3a to IR shifts cells into a quiescent persister state, potentially in large part mediated through abrogating p21 transcriptional repression, leading to downregulation of DREAM target genes.

Finally, we tested whether the transcriptional differences induced by the addition of nutlin-3a to IR, leading to the alleviation of MDM2-mediated repression of p21 transcription, were associated with treatment resistance, as suggested by the increase in persister cell signature scores. We created two gene signatures, consisting of genes either significantly upregulated or downregulated in IR + nutlin-3a versus IR-treated MCF-7 cells (Fig. 6a). Both the upregulated and downregulated gene signatures were associated with IR resistance in cell lines from the Cancer Cell Line Encyclopedia (71) (p = 6.4E-6 and 1.5E-10, respectively; weighted means across multiple primary tumor sites) (Fig. 6e). However, an increase in the upregulated signature was associated with sensitivity to IR in breast and ovarian cancer cell lines. Indeed, in breast cancer patients from the METABRIC cohort (72), the downregulated gene signature was associated with longer recurrence-free (p = 7.1E-4) and overall (p = 0.0055) survival in patients not treated with adjuvant therapy (Supplementary Fig. 8b, c) and shorter recurrence-free and overall survival in patients treated with adjuvant chemo-radiation therapy (p = 0.021 and 0.021, respectively) (Fig. 6f, g). The association between the upregulated signature and recurrence-free or overall survival was borderline significant in patients not treated with adjuvant therapy (p = 0.051 and 0.044, respectively; Supplementary Fig. 8d, e) and was not significant in patients treated with adjuvant chemo-radiation therapy (p = 0.7 and 0.39, respectively; Supplementary Fig. 8f, g). These results are consistent with tumors with lower expression of genes downregulated upon addition of nutlin-3a to IR being less proliferative and more resistant to chemo-radiation therapy. Note that the associations between these signatures and treatment responses does not imply that p21 governs response in these cell lines and tumours. Rather, they suggest that the cell state induced by adding nutlin-3a to IR (which could also be induced by other means, including in TP53-mutant cells) may be resistant to cytotoxic therapy. Taken together, while not definitive, our findings are suggestive of p21-mediated downregulation of genes shifting cells into a quiescent state that is resistant to cytotoxic therapy upon addition of nutlin-3a to IR.

## Discussion

The tumor suppressor gene p53 and its transcriptional target p21 are among the most studied genes in biology. However, despite being the subjects of such extensive investigation, how these genes control cell fate decisions that are vital to successful outcomes of anti-cancer therapies are incompletely understood. Given the role of p21-induced cell cycle arrest in treatment resistance, we aimed to uncover how heterogeneous p53 dynamics are propagated to p21 dynamics using combined single cell microscopy and computational modeling. We found that p21 transcription was far better predicted by the change in p53 than absolute p53 level. Through constructing and comparing stochastic computational models of PR and IFFL network architectures, we determined that p21 transcription is indeed governed by an IFFL. This finding was robust to the type of relationship between p53 levels and p21 promoter activation in the PR model, as neither linear nor Hill functions are able to generate the non-monotonic relationship between p53 levels and p21 transcription observed in the data. We then proposed that MDM2 binding to p53 is responsible for repression of p21 transcription and successfully validated the need for p53-MDM2 binding to explain p21 transcription dynamics. Pharmacological disruption of the IFFL inhibited and reversed the G1-S transition leading to G0 arrest instead of G2/M checkpoint arrest.

Note that this shift in cell fate is specifically related to the perturbation of the network structure and not just the altered p53 dynamics due to reduction in p53 degradation. Given that UV radiation produces similar p53 dynamics to the IR + nutlin-3a treatment used but different cell fates (predominantly apoptosis rather than cell cycle arrest) (6) in the same cell line, we suggest that transcriptional repression of p21 enabled by MDM2 binding to p53 plays an important role in cell fate specification. UV radiation activates ATR, which inhibits MDM2-dependent degradation of p53, but does not alter MDM2 binding to p53 and, hence, the transcriptional activity of p53 (80). Therefore, UV radiation generates similar p53 dynamics to IR + nutlin-3a, but lower p21 induction. However, this difference in cell fates could alternatively be related to different p53 post-translational modifications in response to the different stimuli.

Whilst MDM2-dependent transcriptional repression of p53 target genes has been observed previously (45–50), the reasons why p53 regulates its target genes in this manner has never been understood. We have demonstrated for the first time how the mechanism of p21 transcription can provide potential advantages to cells over simple positive regulation. Specifically, our analyses indicate that the IFFL architecture facilitates rapid cell cycle arrest and increases p21 noise. These findings, in combination with our findings on the influence of p21 protein noise on escape from cell cycle arrest (15), suggest that the IFFL enables a population of cells to rapidly undergo cell cycle arrest following DNA damage and exhibit variability in the timing at which cells exit cell cycle arrest, and become sensitive to subsequent stresses, reducing the probability of population extinction.

Moreover, our combined experimental and mathematical modelling of the transcriptional dynamics of many p53 target genes found that while a small number of genes are regulated by IFFL, most are instead regulated by PR -- or IFFL with only a mild effect of MDM2 on repression of transcription. Such a difference in the modes of transcriptional regulation would enable cells to decode a multiplexed signal encoded in p53 dynamics. In addition to p21, this analysis provided evidence for IFFL regulation of MDM2 transcription. Consistent with this finding, a recent study found that MDM2 transcription is regulated by an IFFL through p53-MDM2 complex binding to the MDM2 promoter and repressing transcription (81). In considering how the expression of different p53 transcriptional target genes could be regulated by these different mechanisms we found, by analyzing TP53 and MDM2 ChIP-seq data, that some p53 transcriptional target genes are bound by p53 alone while others are co-bound by p53 and MDM2, consistent with a previous study (46). This co-binding of p53 and MDM2 occurs at promoters, but not enhancers. This observation provides a potential mechanistic basis for the difference in modes of transcriptional regulation, which warrants further investigation. Our proposed model predicts that disrupting the IFFL through inhibiting the binding between p53 and MDM2 would lead to “crosstalk” between the multiple signals encoded in the p53 dynamics, with transcriptional target genes normally regulated by IFFL to parse the change in p53 instead parsing the absolute level of p53, possibly altering cell fates. This property of the p21 IFFL, inferred from bulk transcriptomics data, would be interesting to explore in more detail in future work by performing single cell measurements of multiple different p53 target genes.

Our findings provide important insights for the design of therapeutic strategies based on p53 activation. Due to the role of MDM2 in transcriptional repression of p21, but not apoptosis genes, inhibiting the p53-MDM2 interaction may steer cell fates towards a temporary G0 arrest, rather than apoptosis, protecting cancer cells from cytotoxic therapies targeting proliferating cells. In support of this assertion, p53 induction by nutlin-3a predominantly leads to cell cycle arrest rather than apoptosis in many cell lines (82–84) and protects melanoma cell lines and patient-derived xenografts from mitotic inhibitor-induced DNA damage in a p21-dependent manner (85). MDM2 antagonists have also been shown to inhibit the senescence-associated secretory phenotype and permanent cell cycle arrest (86). This effect could also be detrimental to successful cancer therapy as induction of tumor cell senescence can increase survival, at least in certain contexts. Given previous work showing that IR-induced senescence results from p53 activation in the G2-phase of the cell cycle (87), our finding that concurrent nutlin-3a and IR administration leads to cells transitioning into the G0-phase of the cell cycle provides a possible explanation for this phenomenon. Alternative explanations include the negative regulation of p53 on mTOR signaling (88). Our findings that nutlin-3a can steer cells into a therapy-resistant quiescent state, together with observations that targeting MDM2 protects a range of p53 WT cancer cells from multiple chemotherapies (85, 89, 90), caution against the use of MDM2 antagonists concurrently with treatments targeting proliferating cells; a strategy currently being investigated in clinical trials (e.g. NCT03107780, NCT05376800, NCT03554707, NCT03217266). Alternative strategies for activating p53 in tumor cells, such as through inhibiting Wip1, may be more likely to cause apoptosis, and hence be superior, to blocking the interaction between p53 and MDM2, which may predominantly result in quiescence.

Our findings indicate that nutlin-3a-induced quiescence could instead be exploited in the context of treating p53-mutant tumors by protecting normal tissues from apoptosis and senescence, without affecting cancer cells, thus widening the therapeutic window. Normal cell senescence is responsible for cancer therapy-associated side effects and can promote the aggressiveness of neighboring cancer cells (91), an effect which is inhibited by MDM2 anatagonists (86). Therefore, steering cells into a temporary G0 arrest, rather than senescence, could be advantageous. This strategy is supported by murine studies demonstrating associations between p21 expression and radioprotection of normal tissue (92–94) and accomplishing p21-dependent radioprotection of normal gastrointestinal tissue by genetically or pharmacologically targeting the p53-MDM2 interaction (95). Moreover, MDM2 agonists have been shown to protect normal cells from multiple chemotherapies (85, 89, 90, 96) in a p21-dependent manner (85). A similar approach has previously been proposed by using CDK4/6 inhibitors to protect normal tissues from radiation through the induction of quiescence (97, 98). However, successful normal tissue protection is highly sensitive to the treatment administration schedule (99). As such, detailed preclinical studies of the schedule-dependent effects of MDM2 antagonists on normal tissue protection should be performed prior to their clinical evaluation in this context.

Our experiments were performed using MCF-7 cells and so the generality of our findings needs to be ascertained in future investigations. Note that MCF-7 cells rarely undergo apoptosis in response to IR due to caspase-3 deficiency, which could potentially affect cell fate following IR, causing cells to undergo cell cycle arrest rather than apoptosis. However, the fact that we observed transcriptional changes consistent with treatment-tolerant persister cells following IR + nutlin-3a suggests that this treatment would be unlikely to predominantly cause apoptosis in a caspase-3 proficient context. Furthermore, IR + nutlin-3a treatment of a caspase-3-proficient MCF-7 cell line (100) predominantly resulted in cell cycle arrest/senescence and minimal apoptosis in a previous study (6). Beyond MCF-7 cells, MDM2 antagonists predominantly lead to cell cycle arrest rather than apoptosis (82–84), although not universally (101), and protect a range of normal and p53 WT cancer cell lines from multiple cytotoxic chemotherapies (85, 89, 90). Moreover, targeting MDM2 protects gastrointestinal tissue from radiation-induced apoptosis through upregulation of p21 and enhances tissue integrity and survival in mice (95). Thus our findings are unlikely to be specific to MCF-7 cells. While MDM2 has been shown to bind the promoter of p21 and block its transcription in a range of cell lines (H1299, D-A2, SJSA-1, MANCA, A875) (45–50), our analysis of TP53 and MDM2 ChIP-seq data from four additional cell lines does suggest that different p53 target genes may be IFFL-regulated in different cells. Therefore, pharmacologically targeting MDM2 may result in different cell fates in different cell types. The mechanism of cell type-specific TP53-MDM2 binding to different p53 target genes remains to be fully elucidated. Our analyses of TP53, MDM2 and H3K27ac ChIP-seq data (Fig. 4h-m) suggest that MDM2 expression levels, histone modifications, co-factor expression and p53 binding site location may all play a role. Future exploration of the extent to which different p53 target genes are regulated by the IFFL in different cell types could help guide selection of patients whose tumors are most likely to respond to MDM2 agonists and normal tissues most likely to be protected against cytotoxic therpies.

In conclusion, we demonstrated that p21 transcription is governed by an IFFL mediated by MDM2 binding to p53, enabling rapid p21 induction following DNA damage and increasing noise in p21 transcription. Pharmacologically abrogating the IFFL leads to G0 arrest, which may protect cells against treatments that target proliferating cells. These findings have potential implications for clinical trials of MDM2 agonists. In particular, combining inhibitors of the p53-MDM2 interaction with agents causing double strand breaks may be beneficial in the treatment of p53-mutant, rather than WT, tumors and alternative p53 activation approaches may be superior in the context treating of p53 WT tumors. Evaluating the optimal use of this class of drugs in light of our findings warrants further investigation in the future.

## Materials and Methods

### Live cell fluorescence microscopy

Live cell fluorescence microscopy, image analysis and quantification of p53 and p21 proteins and p21 transcription following treatment with IR in MCF-7 cells was performed as previously described (18). In an initial experiment, we treated cells with either 2.5 Gy (n = 245), 5 Gy (n = 311) or 10 Gy (n = 279) of IR and imaged them for 45 h with a 15 minute temporal resolution (Fig. 1, Fig. 2, Supplementary Fig. 1a-p, Supplementary Fig. 2a-h, Supplementary Fig. 6d, e). The first 24 h of imaging data of the cells treated with 10 Gy of IR were previously published (18). In a second experiment (18), cells were treated with 10 Gy of IR and imaged for 15 h with a 2 minute temporal resolution (Supplementary Fig. 1q, Supplementary Fig. 2i-l). In a third experiment (18) to validate the computational model predictions, cells were treated with 10 Gy IR alone (n = 249) or in combination with 10 *μ*M nutlin-3a (n = 251) and imaged for 21 h with either a 15 minute (p53, p21-MS2) or 30 minute (p21) temporal resolution (Fig. 3, Fig. 4c, d, Fig. 5a-f, Supplementary Fig. 4, Supplementary Fig. 5, Supplementary Fig. 6a-c, Supplementary Fig. 7h-j). In a fourth experiment to infer the production and degradation rates of p21, cells were treated with 10 Gy IR and then imaged with a 15 minute temporal resolution with 40 nM siRNA targeting either p21 alone or p53 and p21 added at 20 h (Fig. 5c, d, Supplementary Fig. 7g). In these three experiments, 835, 945, 500 and 345 cells, respectively, were successfully segmented and tracked for the duration of the time course. “Outlier” cells with maximum signal intensity greater than 2 interquartile ranges above the 75^th^ quartile, on the log_10_ scale, were removed. This exclusion criterion led to the removal of 5/835 (all 10 Gy IR-treated cells), 2/945, 1/500 (10 Gy IR-treated cell) and 0/345 of the cells in the first, second, third and fourth experiments, respectively. As the fraction of excluded cells was very small, including these cells would minimally affect the results.

### Predictive modeling of p21 gene state based on absolute and change in p53 protein level

Following the “random telegraph” model of gene transcription, we assumed that the p21 gene could be in one of two states: “ON” and “OFF”. We inferred the dynamics of the p21 gene state for each cell by fitting a 2 state hidden Markov model to the p21-MS2 signal for that cell assuming two states and a Gaussian error distribution (Fig. 1f).

We smoothed the p53 signal using loess smoothing with degree 2 and span 0.1 (Fig. 1d). These parameters were chosen as they were found to successfully remove the spikes in the signal without affecting the p53 oscillations.

Delays between the p53 and p21-MS2 signals may occur for biological reasons, such as the time taken for p53 to bind the promoter and recruit the transcriptional machinery, and technical reasons, for example, differences in the maturation times of the different fluorescent reporters. Such delays could bias models. We, therefore, computed the cross-correlation between the p53 and p21-MS2 time course for each cell and shifted the p53 signal to achieve the maximum correlation between the signals. We placed a constraint on the maximum size of the shift of 2.25 h, half the period of p53 oscillations in human cells, to prevent aligning incorrect p53 pulses with bursts of p21 transcription.

We then fit two different univariable logistic regression models to the binary p21 gene state for each cell. For the first model we used the absolute (shifted) p53 level as the covariate (Fig. 1d, g) and for the second model we used the change in (shifted) p53 (Fig. 1e, h). The predictive performance of the models was assessed using the area under the receiver operating characteristic curve and pairwise comparisons of the predictive performances of the models were performed using paired Wilcoxon tests (Fig. 1i, Supplementary Fig. 1p, q).

### Robustness of predictive modeling of p21 gene state to long-term trends in p53 signal

To remove long-term trends in the p53 signal that were present in IR-treated cells, which could represent technical artifacts rather than representing true p53 protein dynamics, we performed b-spline regression with 4 knots on the smoothed p53 signal (Supplementary Fig. 1n, o) and subtracted the fitted spline from the smoothed p53 signal. This number of knots was selected as it was found to successfully remove the longer-term trend in the signal while retaining the p53 oscillations. We then repeated the logistic regression modeling using the detrended p53 signal (Supplementary Fig. 1p).

### Stochastic modeling of p21 transcription dynamics

As the number of nascent p21 RNA were very small we modeled p21 transcription dynamics as stochastic processes, employing continuous time Markov modeling. We chose to use the p53 data as input into the model rather than explicitly modeling the p53 dynamics for two reasons. Firstly, the dynamics of p53 are complex, being affected by multiple other proteins for which we did not have data. Secondly, it allowed the modeling to be informed by the data, reducing the risk of removing biologically informative dynamics.

In the PR model, the p21 promoter can switch between an “OFF” state (*Poff*) and “ON” state (*Pon*), where the switching to the ON state is dependent on the level of p53. RNA molecules (*RNA*) are produced at the transcription site only when the promoter is in the “ON” state. The positive regulation model is defined by the following reactions with per capita rates:

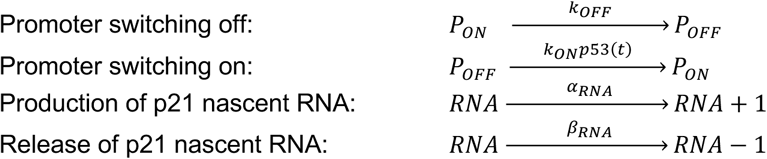

The master equation is given by

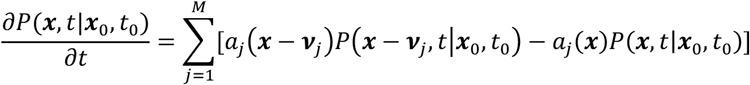

where the state vector for the system is

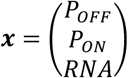

and the set of reactions is described by the state-transition matrix

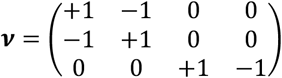

and propensity vector

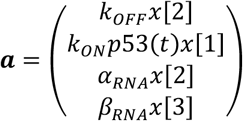

The initial state vector is

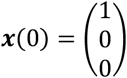

The IFFL model is similar to the PR model, but instead of the promoter switching from the “OFF” to the “ON” state being dependent on the level of p53, it is dependent on the level of p53 above the level of a repressor protein (*R*) whose production is dependent on the level of p53. The model makes the simplifying assumptions of a single step in the production of the repressor (rather than explicitly modeling transcription and translation) and rapid and strong binding of the repressor to p53. The IFFL model is defined by the reactions with the following per capita rates:

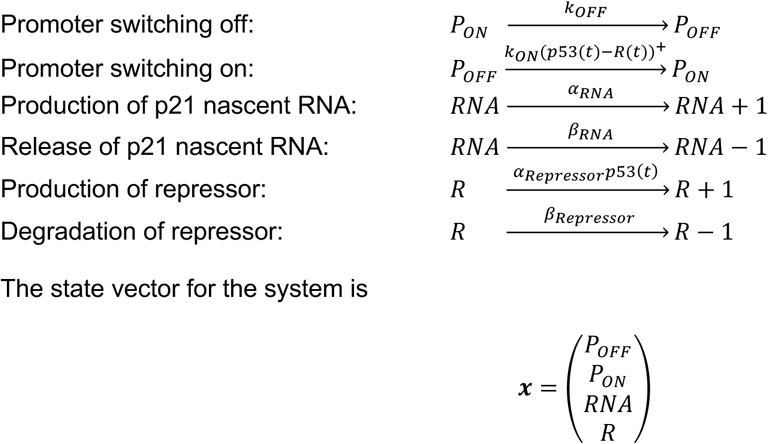

and the set of reactions is described by the state-transition matrix

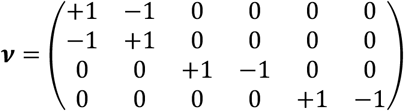

and propensity vector

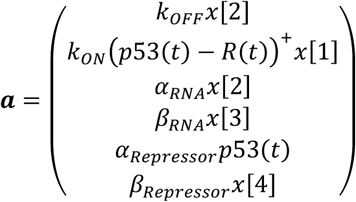

The initial state vector is

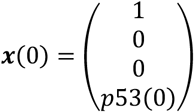

Simulations were performed using the Gillespie stochastic simulation algorithm. For each run of the simulation, we randomly sampled parameter values from the prior distributions, defined in Supplemental Table 1, and simulated p21 transcription dynamics for the same number of cells as in the dataset that the simulations were being compared to. We performed 5,000 simulation runs for each model (following recommendations (102)). To perform the simulations of the time inhomogeneous continuous Markov process we made minor modifications to the ssar R package to increase its speed for our task. This adapted version of the package is available at https://github.com/jamiedean/ssar. Note that we chose to use simple linear relationships for the activation of the p21 promoter by p53 due to the large computational costs of running the simulations and Bayesian inference procedure (see below). We tested the robustness of our model comparison conclusions to this simplification (see below).

### Model comparison by approximate Bayesian computation random forest classification

To select between the two alternative models, we performed model comparison using the approximate Bayesian computation random forest method (102) implemented in the abcrf version 1.9 R package. Briefly, this approach first trains a random forest classifier to predict the identity of the simulation model from the summary statistics extracted from the model simulations. It then applies this classifier to the data to predict the probability with which the data were generated by each model. To formally compare the model simulations to the data we computed the p21 nascent RNA concentration autocorrelation and the cross-correlation between the p53 protein concentration and p21 nascent RNA concentration for all individual cells and took the mean of these functions over all cells, for both the simulations and the data. To account for the maturation time of the p53-tagged fluorescent protein, we introduced a time shift in the p53 protein level time course. The delay was computed as 1/*ρ*, where *ρ* represents the maturation rate of the fluorescent protein. The value of *ρ* was sampled from a prior distribution and so varied between simulations, incorporating uncertainty in the fluorescent reporter maturation. For each value of *ρ* drawn from the prior distribution, the same maturation delay was used for all cells. The auto- and cross-correlation functions were chosen as they incorporate the information from the same single cells over time (103), unlike the moments of distributions over all of the cells at single timepoints. The correlation function values at the first five peaks and troughs of the mean auto-correlation function and the five peaks and troughs closest to a lag of 0 of the mean cross-correlation function were chosen as the summary statistics for Bayesian model comparison. The random forest model used 3,000 trees (a number chosen because the out-of-bag performance had converged with this number of trees) and added linear discriminant analysis to the summary statistics. We only performed model comparison and did not perform parameter inference as this requires a much larger number of parameter sets (100,000 parameter sets recommended (104)), and therefore simulations, which would take several months with our available hardware.

### Robustness of model comparison results to promoter activation dynamics

In order to ensure that our conclusion identifying the IFFL model as the best fit to the experimental data is not an artifact of the simplified form used to model positive regulation, we also implementaed an extended version of the PR model, called PR-Hill Function model. In this model, we define the promoter switching rate from the OFF to the ON state as a Hill function of the p53 level, introducing a nonlinear dependence of activation of p21 transcription by p53. Specifically, the propensity for the promoter switching to the ON state is defined as

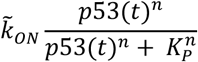

where *k̃_ON_* represents the maximal activation rate, *n* represents the Hill coefficient and

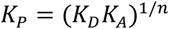

with *K_D_* denoting the dissociation constant of the activator from the DNA promoter site and *K*_A_ denoting the dissociation constant for the cooperative binding reactions for the activator. The overall structure of the model, including the state-transition matrix and state vector, does not change and the only difference from the original PR model is the second entry in the propensity vector, which now include the Hill-type activation function.

### Delay differential equation models of p21 dynamics

Delay differential equation models of p53 and p21 dynamics were developed to perform a mathematically controlled comparison of the effect of the IFFL on the p21 induction rate. The functional form of our model was inspired by two previous models, one of p53 dynamics, incorporating activation of p53 by phosphorylated ATM (pATM) and negative regulation by the p53 transcriptional target genes MDM2 and Wip1 (21), and another of p21 dynamics, incorporating a bistable switch between p21 and CDK2 (15). We added two alternative p21 protein compartments, with p21 production governed either by PR (p21_PR_) or IFFL (p21I_FFL_):

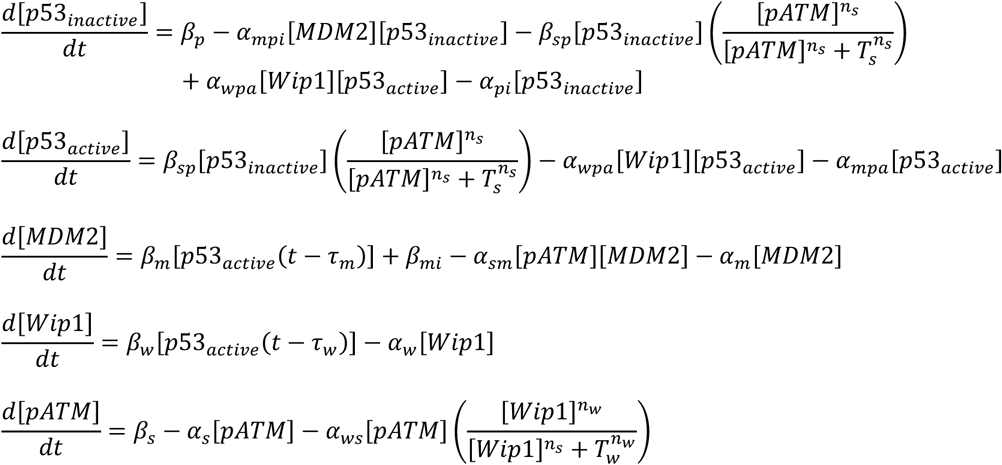

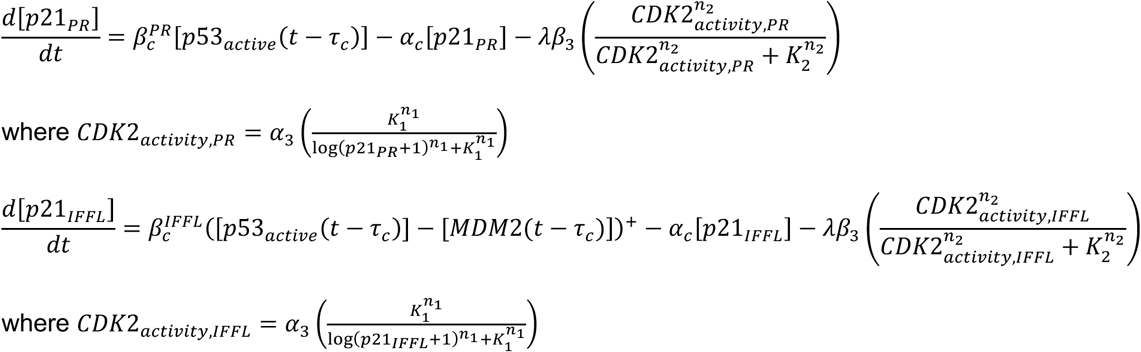

The model parameter values and initial conditions used are given in Supplemental Table 2. We chose the same parameter values and initial conditions as those used in the original publications as these had previously been shown to reproduce the relevant experimental measurements, with the following exceptions: (i) we selected the value for the p21 production rate under positive regulation 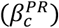 and scaling of the CDK2-dependent p21 degradation (*λ*) to produce similar p21 dynamics as observed in our data; (ii) we selected the p21 production rate under incoherent feedforward loop regulation 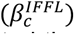 to produce a quasi-steady state level of p21 matching that under positive regulation; (iii) we selected the time delay in p21 production (*τ*_c_) to match that of MDM2. The model was implemented in Julia using the DifferentialEquations package. Note that neither the details of the part of the model that produces the p53 oscillations nor govern p21 degradation affect the relationship between IFFL regulation of p21 production and increased response rate. Alternative mathematical models of p53 dynamics and p21 degradation, demonstrate the same relationship between the mode of p21 regulation and response rate (confirmed when constructing this model).

### Inferring the mode of transcriptional regulation of p53 target genes

Time course RNA-sequencing data and p53 protein levels of MCF-7 cells treated with IR and IR + nutlin-3a were obtained from a previous study (59). Treatment doses and timings are as described previously (6). TP53 transcriptional target genes were defined as those having a p53-dependent mRNA fold change of at least 2, at any time point, and TP53 ChIP-seq peak within 20 kb of the transcription start site. RNA-seq measurements were performed hourly from 0 - 12 h following 10 Gy IR and TP53 ChIP-seq measurements were performed 0 h, 1 h, 2.5 h, 5 h and 7.5 h following 10 Gy IR in MCF-7 cells. The p53 proteins were smoothed using a smoothing spline with 12 knots, a number selected based on successfully capturing the known patterns of p53 dynamics following IR and IR + nutlin-3a.

The dynamics of p53 transcriptional target gene mRNA expression, *M*, were modelled using ordinary differential equation models with transcription governed by either PR or IFFL:

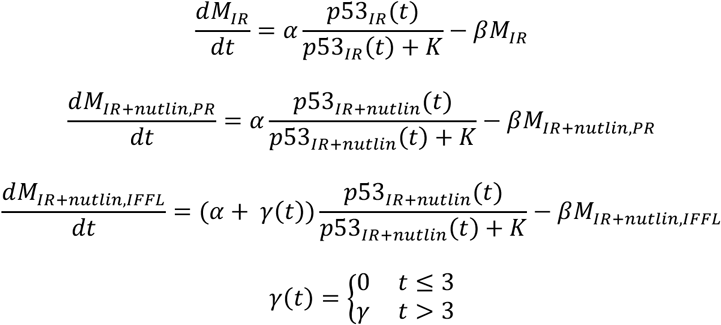

where *α* is the transcription rate, *β* is the degradation rate, *K* is the Michaelis-Menten constant, and *γ* is the increase in transcription rate due to inhibition of transcriptional repression in the presence of nutlin-3a. The time dependence of *γ* is due to nutlin-3a being administered 3 h after IR administration.

mRNA expression dynamics were simulated under both models for 10,000 different combinations of parameter values, with parameters sampled from the prior distributions defined in Supplemental Table 3. Model comparison was performed using approximate Bayesian computation random forest method (102) implemented in the abcrf version 1.9 R package. The mRNA levels at each timepoint under both treatment conditions were used as the summary statistics. For the data, the mean values across the two replicates were used. The random forest model used 3,000 trees (a number selected because the out-of-bag performance had converged with this number of trees) and added linear discriminant analysis to the summary statistics.

### ChIP-seq data analysis

TP53 and MDM2 ChIP-seq data of four cell lines were obtained from a previous study (60). Overlaps between TP53 and MDM2 peaks and distances from the centers of TP53 peaks to transcription start sites were computed using the ChIPpeakAnno version 3.40.0 R package, with genes annotated using the TxDb.Hsapiens.UCSC.hg38.knownGene version 3.20.0 R package. ChIP-seq tracks were plotted using SparK version 3.0 (105). The analysis of distances between TP53 peaks and transcription start sites was not performed for the HCT116 and U2OS cell lines as the number of overlapping TP53 and MDM2 peaks was too small (3 and 4 overlapping TP53 and MDM2 peaks, respectively) to perform a meaningful statistical analysis.

### Theoretical modeling for inference of p21 production and degradation rates

To infer p21 production and degradation rates from time-lapse microscopy data, we begin by assuming that p21 protein concentration, [*p*21], is described by the ordinary differential equation

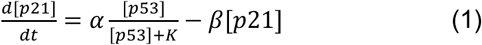

where *α* is rate of production of the p21 protein, *K* is the Michaelis-Menten constant and *β* is the degradation rate of the p21 protein. At quasi-steady state, ^P[(G2]^ ≈ 0, and equation (1) can be rearranged to give

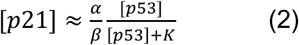

If [*p*53] ≫ *K*, then

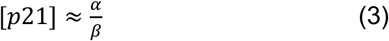

Therefore, given the assumptions of this model, for sufficiently high levels of p53, plotting [*p*21] against [p53] would give rise to a straight line with slope of 0 and y-intercept of 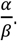 If this theoretical insight is correct, and assuming that p21 degradation rate is unaffected by nutlin-3a, this argument would enable the relative difference in the p21 production rate between cells treated with IR alone and IR + nutlin-3a to be inferred. Measuring the degradation rate of p21 (method described below) enables the absolute p21 production rates in the absence and presence of nutlin-3a to be inferred by

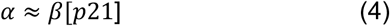

As p21 has two different degradation rates: rapid degradation in S-phase of the cell cycle and slow degradation in the remainder of the cell cycle (27), Gaussian mixture models with 2 components were fit to the [*p21*] data for each treatment condition. For each treatment condition, the p21 production rate was inferred separately for each of the two mixture components and the inverse variance weighted mean was used to combine these estimates into a single production rate estimate.

To satisfy the quasi-steady state assumption, we restricted our analysis to the final 2.5 h of the experiment and to cells for which the magnitude of the slope of a linear regression model fit to the final 2.5 h of the p21 time course data was less than 0.05 a.u.h^-1^.

To predict the effect of abrogating MDM2-mediated transcriptional repression without affecting p53 degradation, and therefore p53 levels, in S-phase cells (rapid p21 degradation), we multiplied the p21 levels of IR-treated cells in the mixture component with lower p21 levels by the ratio of production rates inferred for IR + nutlin-3a- and IR-treated cells (Fig. 5f). To predict the effect of abrogating MDM2-mediated p53 degradation without affecting transcriptional repression, we added the difference in the mean p53 levels between IR + nutlin-3a- and IR-treated cells to the p53 levels of IR-treated cells in the mixture component with lower p21 levels (Supplementary Fig. 7h). In both cases we computed standard errors by propagating errors in the rate and protein steady state level estimates.

### Measurement and inference of p21 degradation rates

MCF-7 cells were treated with 10 Gy IR and then imaged with a 15 minute temporal resolution with 40 nM siRNA targeting either p21 alone or p53 and p21 added at 20 h. Piecewise linear regression models were fit to the log-transformed p21 protein signal versus time data for each cell individually, using the data from 27.5 h onwards only (to allow time sufficient time for p21 protein production to be inhibited), with the dpseg version 0.1.1 R package. The slopes of the first two lines were taken to be the low and high degradation rates for each cell. A Gaussian mixture model with 2 components was fit to the distribution of inferred degradation rates across all cells (combining cells from both datasets as there was the degradation dynamics were the same under both conditions) using the mclust version 6.1.1 R package.

### RNA-sequencing data analysis

Raw RNA-seq time course data of MCF-7 cells sequenced from 3 h to 12 h in 1 h intervals following treatment with IR and IR + nutlin-3a (59) were downloaded from the Gene Expression Omnibus (GSE100099). Expression of transcripts was quantified using Salmon version 1.1.0 (106) with GENCODE release 33 (107) Homo sapiens GRCh38 for the reference transcriptome annotation. Differential expression analysis was performed to test for differences in gene expression over time between treatments using the likelihood ratio test, with expression ∼ time + replicate + treatment + treatment:time as the full model and expression ∼ time + replicate + treatment as the reduced model, in the DESeq2 version 1.46.0 R package.

### Transcription factor enrichment analysis

Transcription factor enrichment analysis was performed using the TFEA.ChIP version 1.26.0 R package (108) with the ReMap2022+EnsTSS+CellTypeEnh.Rdata database accessed from https://github.com/LauraPS1/TFEA.ChIP_downloads/tree/master/R%20Databases. Over-representation analysis of transcriptional regulators for significantly upregulated and downregulated genes was performed. Upregulated and downregulated genes were defined as those with a differential expression adjusted p-value less than 0.05 and a positive or negative estimates of differences in gene expression over time, respectively. Genes with a differential expression adjusted p-value greater than 0.5 were selected as control genes.

### Transcriptional signatures

Transcriptional signature scores were calculated with gene set variance analysis, implemented in the GSVA version 2.0.7 R package. Cell cycle phase (65), nuclear size (109) and persister cell signatures^55–62^ were obtained from the referenced studies.

Gene expression signatures were created for genes significantly upregulated and downregulated in IR + nutlin-3a- versus IR-treated cells with an adjusted p-value of less than 0.05 and effect size of magnitude greater than 0.5. The associations between the upregulated and downregulated gene expression signatures and cell line radiation response were measured by fitting linear regression models with area under the radiation dose response curve as the outcome variable and the gene set variance analysis signature scores as the covariate, using data from the RadioGx version 2.10.0 R package (71). Separate regression models were fit for cell lines from different primary disease sites, for disease sites with at least 10 cell lines, and an inverse variance weighted mean coefficient across all of the primary disease sites calculated (Fig. 6e).

The associations between the upregulated and downregulated gene expression signatures and breast cancer patient recurrence-free and overall survival were measured by fitting Cox proportional hazards regression models with the gene set variance analysis signature scores as the covariates, using data from the METABRIC cohort (72) (Fig. 6f, g and Supplementary Fig. 8b-g). Patients treated with hormone therapy (n = 1216) were excluded as hormone therapy could potentially confound inference of cytotoxic therapy response, leaving 311 patients receiving no adjuvant therapy and 173 patients treated with radiation and chemotherapy.

Note that cell lines and patients with TP53 mutations were not excluded from these analyses as the purpose was to assess the association between the transcriptional state induced by the addition of nutlin-3a to IR and treatment response. Such a state could be induced by other means, including in a p21-indpendent manner and in TP53 mutant cells, so cell lines and tumors were not removed on the basis of any molecular features.

## Materials availability

This study did not generate new unique reagents.

## Data Availability

Data are available at https://osf.io/5w4gs/ (DOI: 10.17605/OSF.IO/5W4GS).

## Code Availability

Code is available at https://github.com/jamiedean/p21-iffl-regulation.

## Acknowledgements

We thank members of the Michor and Lahav labs for discussion and comments on the manuscript. This project was supported by funding from the Dana-Farber Cancer Institute Physical Science Oncology Center (NCI U54CA193461 to F.M.), the National Institutes of Health grant R35GM139572 (G.L.) and the Ludwig Center at Harvard (G.L., F.M.). This work was supported by the Radiation Research Unit at the Cancer Research UK City of London Centre Award [C7893/A28990].

## Author Contributions

Conceptualization, J.D., J.R., G.L. and F.M.; Methodology, J.D., J.R., S.B.; Software, J.D., J.R., S.B.; Formal Analysis, J.D.; Investigation, J.D.; Writing – Original Draft, J.D.; Writing – Review & Editing, J.D., J.R., S.B., M.T., A.J., G.L. and F.M.; Visualization, J.D.; Supervision, G.L. and F.M.; Funding Acquisition, G.L. and F.M.; Resources, J.R., M.T., A.J. and G.L.

## Declaration of Interests

The authors declare the following competing interests: F.M. is a cofounder and advisor to Harbinger Health, an advisor to Zephyr AI, and serves on the board of directors of Recursion Pharmaceuticals. F.M. declares that none of these relationships are directly or indirectly related to the content of this manuscript.

## Notes

https://osf.io/5w4gs/

https://github.com/jamiedean/p21-iffl-regulation

### Summary of Updates

Figures 1, 4, 5 and 6 revised; Supplementary Table 1 revised; Supplementary Figures 3 and 5 added; Supplementary Figures 1, 4, 6 and 7 revised; text revised to include new methods and results, and clarify methodological details and interpretations of results; author added.

