## Supplementary Materials for "Functional consequences of a p53-MDM2-p21 incoherent feedforward loop"

**Supplementary Table 1: Prior distributions for the stochastic models of p21 transcription dynamics.**

| **Parameter** | **Prior Distribution** | **Unit** | **Justification** |
| --- | --- | --- | --- |
| $k_{OFF}$ | $\mathcal{N}_{T}\left( 500, 500, 0, \infty;x \right)=\left\{ \begin{aligned} \mathcal{N}\left( 500, 500;x \right) \\ 0 \end{aligned}{\mathrm{if} x\geq0 \atop\mathrm{if} x<0} \right.$ | h^-1^ | Previous measurements in mammalian cells^1^. Should be substantially larger than $k_{ON}$ to get bursts of transcription. |
| $k_{ON}$ | $\mathcal{N}_{T}\left( 1, 10, 0, \infty;x \right)=\left\{ \begin{aligned} \mathcal{N}\left( 1, 10;x \right) \\ 0 \end{aligned}{\mathrm{if} x\geq0 \atop\mathrm{if} x<0} \right.$ | h^-1^ | Previous measurements in mammalian cells^1^. |
| $\alpha_{RNA}$ | $\mathcal{N}_{T}\left( 250, 250, 0, \infty;x \right)=\left\{ \begin{aligned} \mathcal{N}\left( 250, 250;x \right) \\ 0 \end{aligned}{\mathrm{if} x\geq0 \atop\mathrm{if} x<0} \right.$ | mRNA h^-1^ | Previous measurements in mammalian cells^1^. |
| $\beta_{RNA}$ | $\mathcal{N}_{T}\left( 7, 10, 0, \infty;x \right)=\left\{ \begin{aligned} \mathcal{N}\left( 7, 10;x \right) \\ 0 \end{aligned}{\mathrm{if} x\geq0 \atop\mathrm{if} x<0} \right.$ | h^-1^ | Previous measurements in mammalian cells^1^. |
| $\alpha_{Repressor}$ | $\mathcal{N}_{T}\left( 0.15, 1, 0, \infty;x \right)=\left\{ \begin{aligned} \mathcal{N}\left( 0.15, 1;x \right) \\ 0 \end{aligned}{\mathrm{if} x\geq0 \atop\mathrm{if} x<0} \right.$ | proteins h^-1^ | Should allow Repressor to track p53 dynamics. |
| $\beta_{Repressor}$ | $\mathcal{N}_{T}\left( 0.15,1, 0, \infty;x \right)=\left\{ \begin{aligned} \mathcal{N}\left( 0.15, 1;x \right) \\ 0 \end{aligned}{\mathrm{if} x\geq0 \atop\mathrm{if} x<0} \right.$ | h^-1^ | Should allow Repressor to track p53 dynamics. |
| $\rho$ | $\mathcal{N}_{T}\left( 1.2, 1.2, 0.15, \infty;x \right)=\left\{ \begin{aligned} \mathcal{N}\left( 1.2, 1.2;x \right) \\ 0 \end{aligned}{\mathrm{if} x\geq0.15 \atop\mathrm{if} x<0.15} \right.$ | h^-1^ | Previous measurements in multiple cell types^2^. |
| $n$ | $\mathcal{N}_{T}\left( 3, 3, 0, \infty;x \right) =\left\{ \begin{aligned} \mathcal{N}\left( 3, 3;x \right) \\ 0 \end{aligned}{\mathrm{if} x\geq0 \atop\mathrm{if} x<0} \right.$ | No units | Captures the observed range of Hill coefficients (1–10) across biological systems^3,4^. |
| $K_{P}$ | $\mathcal{N}_{T}\left( 500, 500, 0, \infty;x \right) =\left\{ \begin{aligned} \mathcal{N}\left( 500, 500;x \right) \\ 0 \end{aligned}{\mathrm{if} x\geq0 \atop\mathrm{if} x<0} \right.$ | proteins | Previous experiments^5^. |

$\mathcal{N}$ – normal distribution; $\mathcal{N}_{T}$ – truncated normal distribution.

**Supplementary Table 2: Delay differential equation model parameter values and initial conditions.**

| **Parameter/Initial Condition** | **Description** | **Value** | **Unit** |
| --- | --- | --- | --- |
| $\beta_{p}$ | p53_inactive_ production rate | 0.9 | C_S_ h^-1^ |
| $\beta_{sp}$ | Saturating production rate of p53_active_ | 10 | h^-1^ |
| $\beta_{m}$ | p53-dependent MDM2 production rate | 0.9 | h^-1^ |
| $\beta_{mi}$ | p53-independent MDM2 production rate | 0.2 | C_S_ h^-1^ |
| $\beta_{w}$ | Wip1 production rate | 0.25 | h^-1^ |
| $\beta_{s}$ | pATM production rate | 10 | C_S_ h^-1^ |
| $\beta_{c}^{PR}$ | p21 production rate with PR | 0.8 | h^-1^ |
| $\beta_{c}^{IFFL}$ | p21 production rate with IFFL | 2.55 | h^-1^ |
| $\alpha_{mpi}$ | MDM2-dependent p53_inactive_ degradation rate | 5 | C_S_^-1^ h^-1^ |
| $\alpha_{wpa}$ | Wip1-dependent p53_active_ inactivation rate | 0.14 | C_S_^-1^ h^-1^ |
| $\alpha_{pi}$ | MDM2-independent p53_inactive_ degradation rate | 2 | h^-1^ |
| $\alpha_{mpa}$ | MDM2-dependent p53_active_ degradation rate | 1.4 | h^-1^ |
| $\alpha_{sm}$ | pATM-dependent MDM2 inactivation rate | 0.5 | C_S_^-1^ h^-1^ |
| $\alpha_{m}$ | MDM2 degradation rate | 1 | h^-1^ |
| $\alpha_{w}$ | Wip1 degradation rate | 0.7 | h^-1^ |
| $\alpha_{s}$ | Wip1-independent pATM degradation rate | 7.5 | h^-1^ |
| $\alpha_{ws}$ | Saturating Wip1-dependent degradation rate | 50 | h^-1^ |
| $\alpha_{c}$ | p21 degradation rate | 0.08 | h^-1^ |
| $n_{s}$ | Hill coefficient of p53_active_ production by pATM | 4 | No units |
| $T_{s}$ | pATM concentration for half-maximal p53 production | 1 | C_S_ |
| $n_{w}$ | Hill coefficient of signal degradation by Wip1 | 4 | No units |
| $T_{w}$ | Wip1 concentration for half-maximal pATM degradation | 0.2 | C_S_ |
| $\tau_{m}$ | Time delay in MDM2 production | 0.7 | h |
| $\tau_{w}$ | Time delay in Wip1 production | 1.25 | h |
| $\tau_{c}$ | Time delay in p21 production | 0.7 | h |
| $\alpha_{3}$ | Maximum CDK2 activity | 1.084 | No units |
| $n_{1}$ | Hill coefficient of p21-dependent CDK2 inhibition | 5.333 | No units |
| $K_{1}$ | p21 concentration for half-maximal p21-dependent CDK2 inhibition | 4.874 | C_S_ |
| $\lambda$ | Scaling of CDK2-dependent p21 degradation | 0.75 | No units |
| $\beta_{3}$ | Maxmimum rate of CDK2-induced p21 degradation rate | 1.6 | h^-1^ |
| $n_{2}$ | Hill coefficient of CDK2-dependent p21 degradation rate | 2.774 | No units |
| $K_{2}$ | CDK2 activity for half-maximal CDK2-dependent p21 degradation rate | 0.917 | C_S_ |
| $\left[ {p53}_{inactive} \right](t=0)$ | Initial p53_inactive_ concentration | 0.3 | C_S_ |
| $\left[ {p53}_{active} \right](t=0)$ | Initial p53_active_ concentration | 0 | C_S_ |
| $\left[ MDM2 \right](t=0)$ | Initial MDM2 concentration | 0.2 | C_S_ |
| $\left[ Wip1 \right](t=0)$ | Initial Wip1 concentration | 0 | C_S_ |
| $\left[ pATM \right](t=0)$ | Initial phosphorylated ATM concentration | 0 | C_S_ |
| $\left[ {p21}_{PR} \right](t=0)$ | Initial p21 concentration with PR | 0 | C_S_ |
| $\left[ {p21}_{IFFL} \right](t=0)$ | Initial p21 concentration with IFFL | 0 | C_S_ |

C_S_ – simulated arbitrary concentration units.

**Supplementary Table 3: Prior distributions for the ordinary differential equation models of p53 transcriptional target gene mRNA dynamics.**

| **Parameter** | **Prior Distribution** | **Unit** | **Justification** |
| --- | --- | --- | --- |
| $\alpha$ | $\mathcal{N}_{T}\left( 1, 5, 0, \infty;x \right)=\left\{ \begin{aligned} \mathcal{N}\left( 1, 5;x \right) \\ 0 \end{aligned}{\mathrm{if} x\geq0 \atop\mathrm{if} x<0} \right.$ | mRNA h^-1^ | Previous measurements in mammalian cells^6^. |
| $\beta$ | $\mathcal{N}_{T}\left( 0, 0.3, 0, \infty;x \right)=\left\{ \begin{aligned} \mathcal{N}\left( 0, 0.3;x \right) \\ 0 \end{aligned}{\mathrm{if} x\geq0 \atop\mathrm{if} x<0} \right.$ | h^-1^ | Previous measurements in mammalian cells^6^. |
| $K$ | $\mathcal{N}_{T}\left( 0.1, 10, 0, \infty;x \right)=\left\{ \begin{aligned} \mathcal{N}\left( 0.1, 10;x \right) \\ 0 \end{aligned}{\mathrm{if} x\geq0 \atop\mathrm{if} x<0} \right.$ | a.u. | Produces production rates consistent with those measured in mammalian cells. |
| $\gamma$ | $\mathcal{N}_{T}\left( 1, 10, 0, \infty;x \right)=\left\{ \begin{aligned} \mathcal{N}\left( 1, 10;x \right) \\ 0 \end{aligned}{\mathrm{if} x\geq0 \atop\mathrm{if} x<0} \right.$ | mRNA h^-1^ | Produces production rates consistent with those measured in mammalian cells. |

$\mathcal{N}$ – normal distribution; $\mathcal{N}_{T}$ – truncated normal distribution.

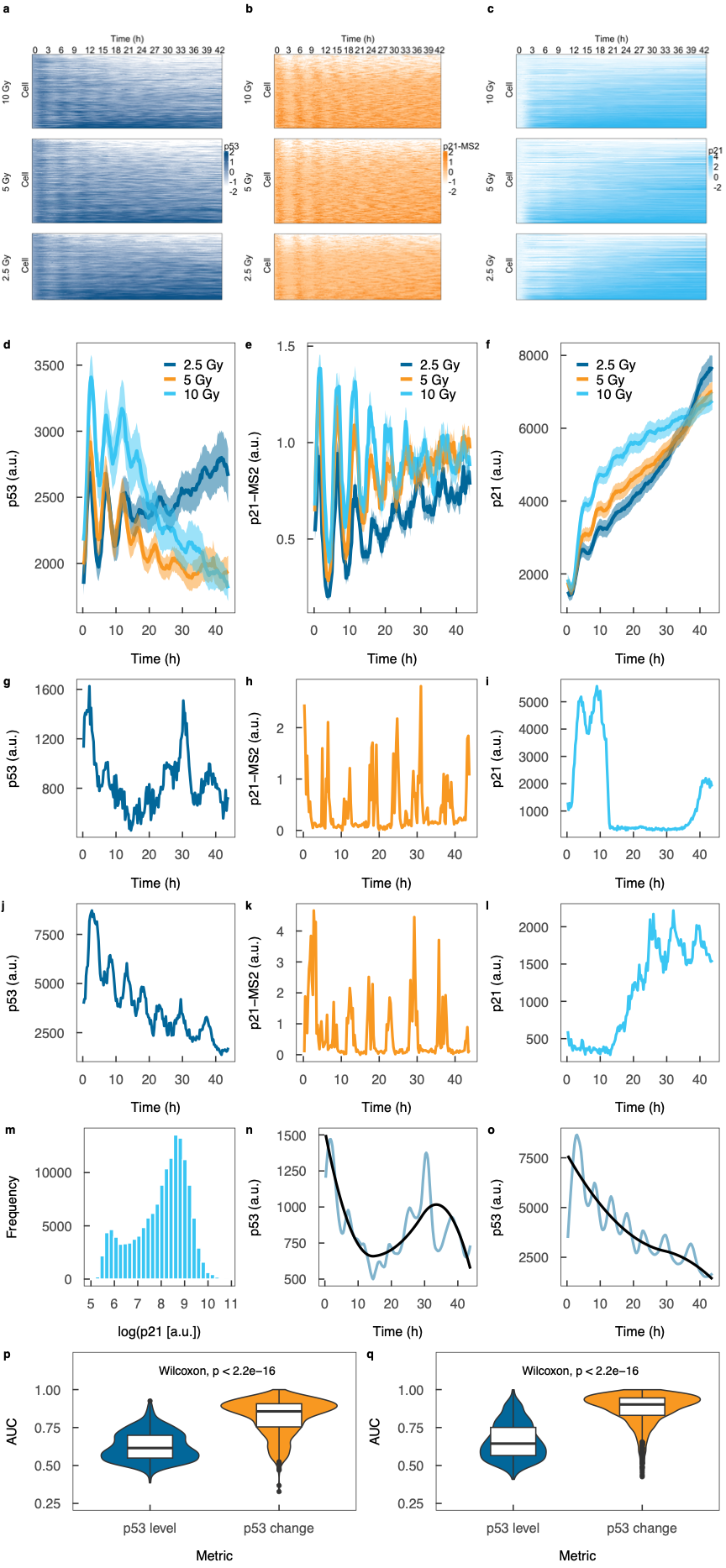

**Supplementary Figure 1: Transcription of p21 is dependent on the change in p53 rather than its absolute level. (a)** Single cell p53 protein dynamics following IR administration. **(b)** Single cell p21-MS2 dynamics following IR administration. **(c)** Single cell p21 protein dynamics following IR administration. The number of cells treated with 10 Gy, 5 Gy and 2.5 Gy IR were 274, 311 and 245, respectively. The color scales show the log_2_ fold change in the fluorescence level and are truncated at the 5^th^ and 95^th^ percentiles. **(d)** Mean p53 protein dynamics following IR administration. **(e)** Mean p21-MS2 dynamics following IR administration. **(f)** Mean p21 protein dynamics following IR administration. The shaded areas indicate the standard errors in the means. **(g)** p53 protein dynamics for a representative cell with a decreasing and increasing trend. **(h)** p21-MS2 dynamics for the same representative cell as in (g). **(i)** p21 protein dynamics for the same representative cell as in (g). **(j)** p53 protein dynamics for a representative cell with a decreasing trend. **(k)** p21-MS2 dynamics for the same representative cell as in (j). **(l)** p21 protein dynamics for the same representative cell as in (j). **(m)** Fit of the trend (black line) to the smoothed p53 protein dynamics (blue line) for the same representative cell as in (g). **(n)** Fit of the trend (black line) to the smoothed p53 protein dynamics (blue line) for the same representative cell as in (j). **(o)** Distribution of p21 protein levels across all cells (n = 834) and timepoints (n = 175), exhibiting a bimodal distribution. **(p)** Predictive performance of the logistic regression models based on the absolute p53 expression level and change in p53 expression level, following detrending of the p53 time course data, for all of the cells in the longer time course experiment (n = 830). **(q)** Predictive performance of the logistic regression models based on the absolute p53 expression level and change in p53 expression level for the shorter time course experiment (n = 945)**.** AUC = area under receiver operating characteristic curve.

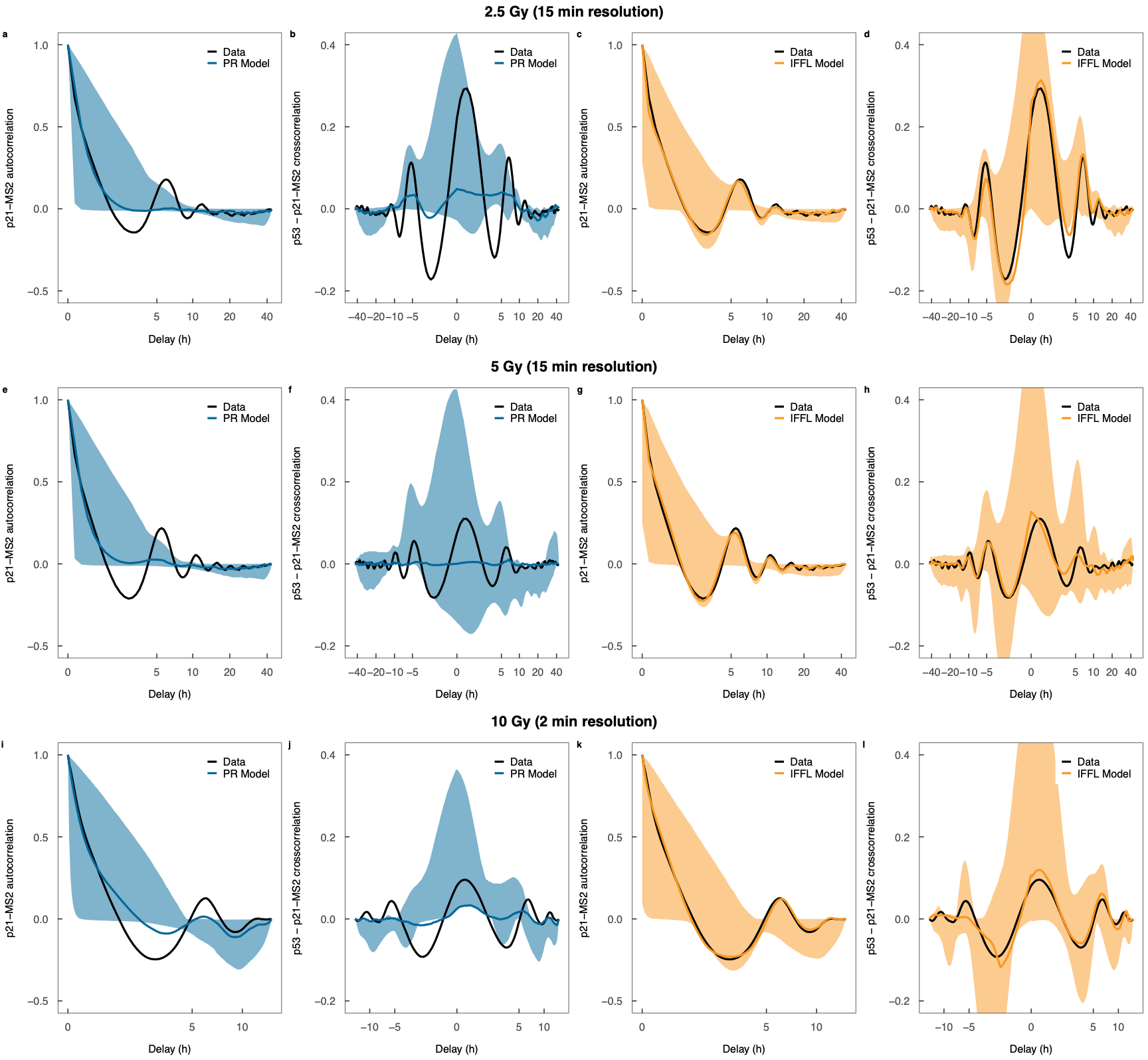

**Supplementary Figure 2: Transcription of p21 is governed by an incoherent feedforward loop that enables p53 change detection. (a)** Mean p21-MS2 autocorrelation function for 2.5 Gy IR-treated cells for the PR model. **(b)** Mean p53-p21-MS2 cross-correlation function for 2.5 Gy IR-treated cells for the PR model. **(c)** Mean p21-MS2 autocorrelation function for 2.5 Gy IR-treated cells for the IFFL model. **(d)** Mean p53-p21-MS2 cross-correlation function for 2.5 Gy IR-treated cells for the IFFL model. **(e)** Mean p21-MS2 autocorrelation function for 2.5 Gy IR-treated cells for the PR model. **(f)** Mean p53-p21-MS2 cross-correlation function for 5 Gy IR-treated cells for the PR model. **(g)** Mean p21-MS2 autocorrelation function for 5 Gy IR-treated cells for the IFFL model. **(h)** Mean p53-p21-MS2 cross-correlation function for 5 Gy IR-treated cells for the IFFL model. **(i)** Mean p21-MS2 autocorrelation function for cells from the short time course experiment (10 Gy IR) for the PR model. **(j)** Mean p53-p21-MS2 cross-correlation function for cells from the short time course experiment (10 Gy IR) for the PR model. **(k)** Mean p21-MS2 autocorrelation function for cells from the short time course experiment (10 Gy IR) for the IFFL model. **(l)** Mean p53-p21-MS2 cross-correlation function for cells from the short time course experiment (10 Gy IR) for the IFFL model. In all panels the colored lines represent the correlation functions from the simulations that are closest to the correlation functions from the data and the shaded areas represent the 95 percentile confidence intervals of the correlation functions from the simulations. Correlation functions are the means of 245, 311 and 945 cells treated with 2.5 Gy, 5 Gy and 10 Gy (short time course experiment) IR, respectively.

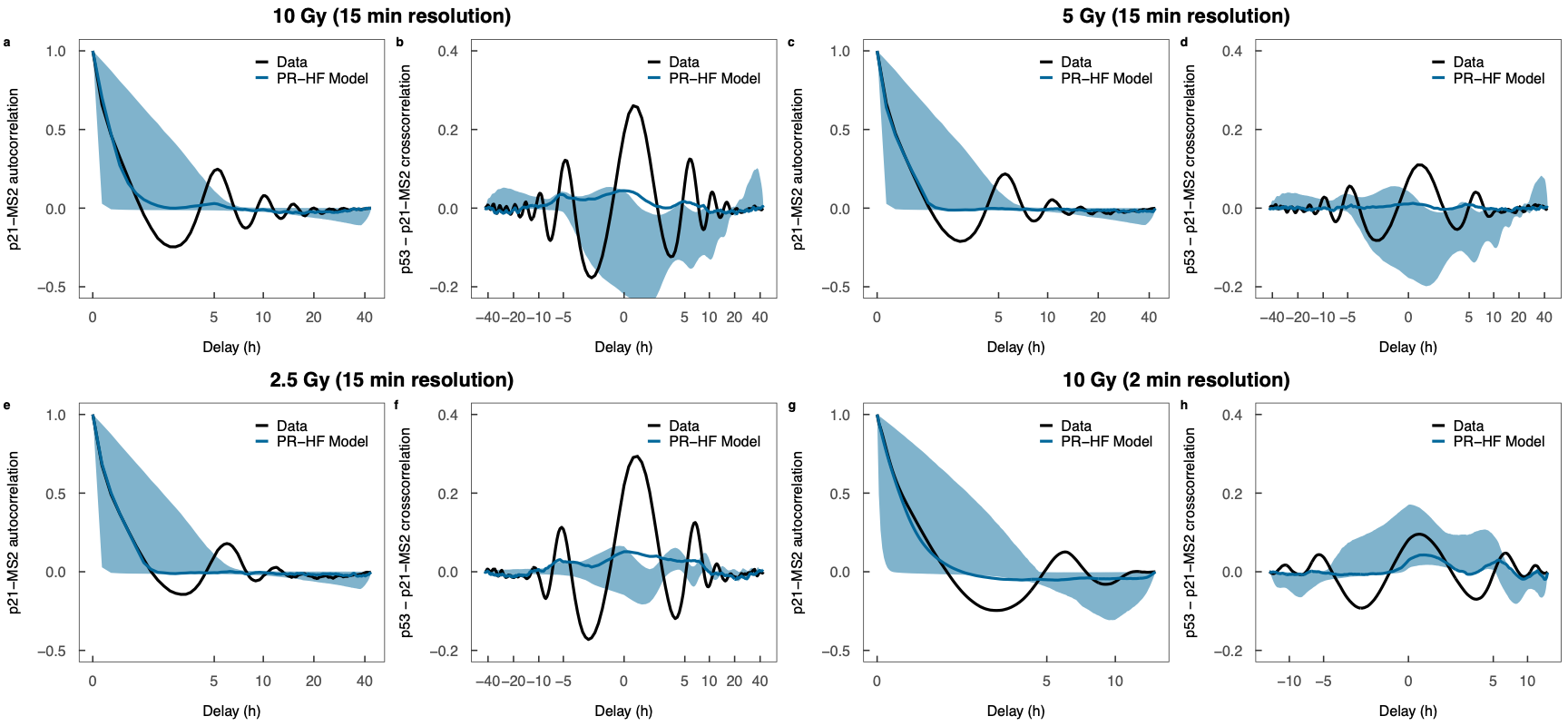

**Supplementary Figure 3: Comparison of experimental and simulated correlation functions when using the PR-Hill Function model.** **(a)** Mean p21-MS2 autocorrelation function for the PR-Hill Function model for cells treated with 10 Gy IR and imaged with a 15 minute temporal resolution. **(b)** Mean p53-p21-MS2 cross-correlation function for the PR-Hill Function model for cells treated with 10 Gy IR and imaged with a 15 minute temporal resolution. **(c)** Mean p21-MS2 autocorrelation function for the PR-Hill Function model for cells treated with 5 Gy IR and imaged with a 15 minute temporal resolution. **(d)** Mean p53-p21-MS2 cross-correlation function for the PR-Hill Function model for cells treated with 5 Gy IR and imaged with a 15 minute temporal resolution. **(e)** Mean p21-MS2 autocorrelation function for the PR-Hill Function model for cells treated with 2.5 Gy IR and imaged with a 15 minute temporal resolution. **(f)** Mean p53-p21-MS2 cross-correlation function for the PR-Hill Function model for cells treated with 2.5 Gy IR and imaged with a 15 minute temporal resolution. **(g)** Mean p21-MS2 autocorrelation function for the PR-Hill Function model for cells treated with 10 Gy IR and imaged with a 2 minute temporal resolution. **(h)** Mean p53-p21-MS2 cross-correlation function for the PR-Hill Function model for cells treated with 10 Gy IR and imaged with a 2 minute temporal resolution. In al panels, the colored lines represent the correlation functions from the simulations that are closest to the correlation functions from the data and the shaded areas represent the 95 percentile confidence intervals of the correlation functions from the simulations. Correlation functions are the means of 274, 311, 245 and 945 cells treated with 10 Gy, 5 Gy, 2.5 Gy and 10 Gy (short time course experiment) IR, respectively.

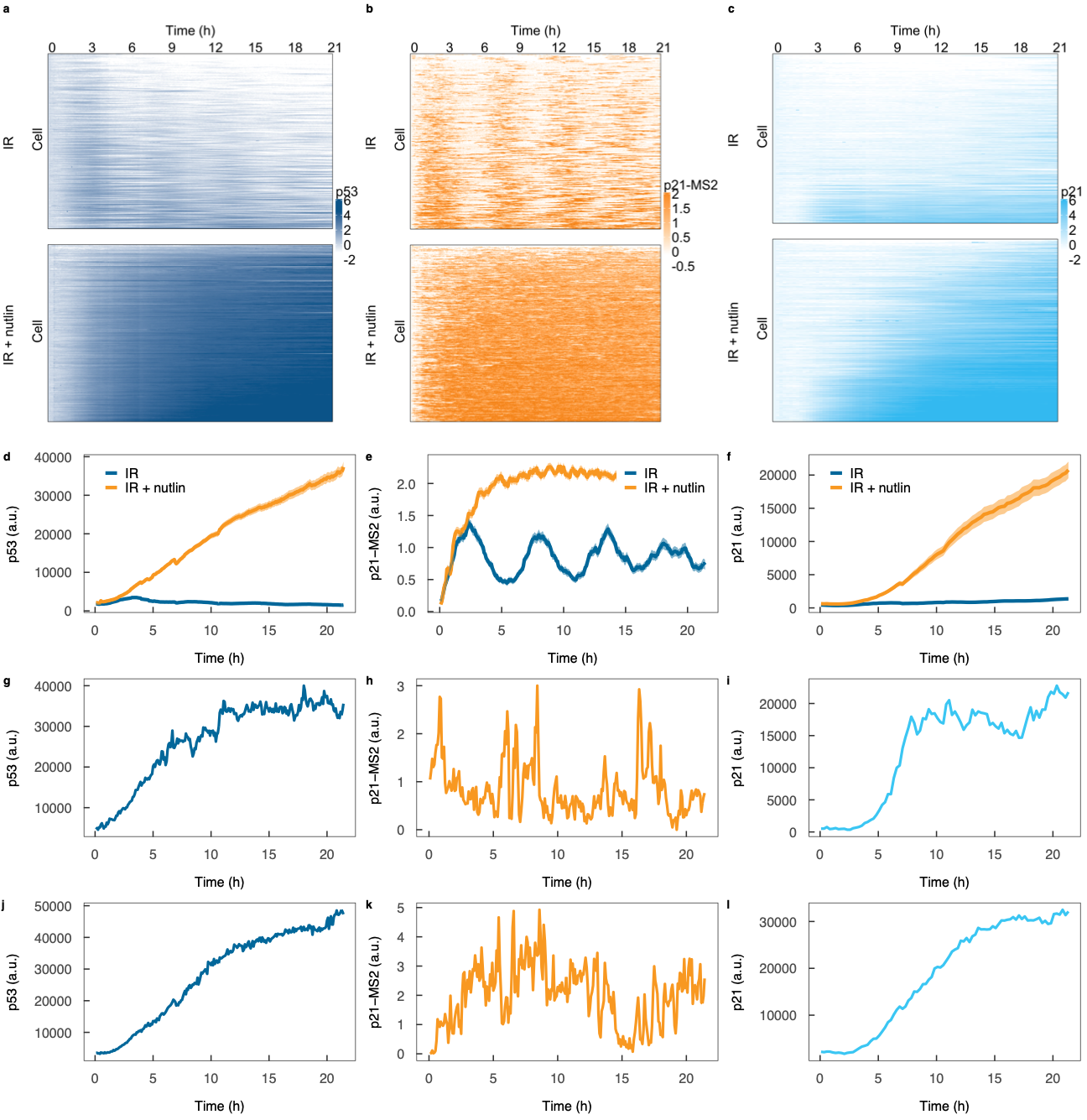

**Supplementary Figure 4: Inhibiting the p53-MDM2 interaction abrogates p53 change-dependent p21 transcription. (a)** Single cell p53 dynamics following 10 Gy IR and 10 Gy IR + 10 $\mu$M nutlin-3a administration. **(b)** Single cell p21-MS2 dynamics following 10 Gy IR and 10 Gy IR + 10 $\mu$M nutlin-3a administration. **(c)** Single cell p21 dynamics following 10 Gy IR and 10 $\mu$M IR + nutlin-3a administration. The number of cells treated with 10 Gy IR and 10 Gy IR + 10 $\mu$M nutlin-3a were 248 and 251, respectively. The color scales show the log_2_ fold change in the fluorescence level and are truncated at the 5^th^ and 95^th^ percentiles. **(d)** Mean p53 protein dynamics following treatment administration. **(e)** Mean p21-MS2 dynamics following treatment administration. **(f)** Mean p21 protein dynamics following treatment administration. The shaded areas indicate the standard errors in the means. **(g)** p53 protein dynamics for a representative cell treated with 10 Gy IR + 10 $\mu$M nutlin-3a. **(h)** p21 transcription dynamics for the same representative cell as in (g). **(i)** p21 protein dynamics for the same representative cell as in (g). **(j)** p53 protein dynamics for another representative cell treated with 10 Gy IR + 10 $\mu$M nutlin-3a (different to the cell in ((g) – (i)). **(k)** p21 transcription dynamics for the same representative cell as in (j). **(l)** p21 protein dynamics for the same representative cell as in (j).

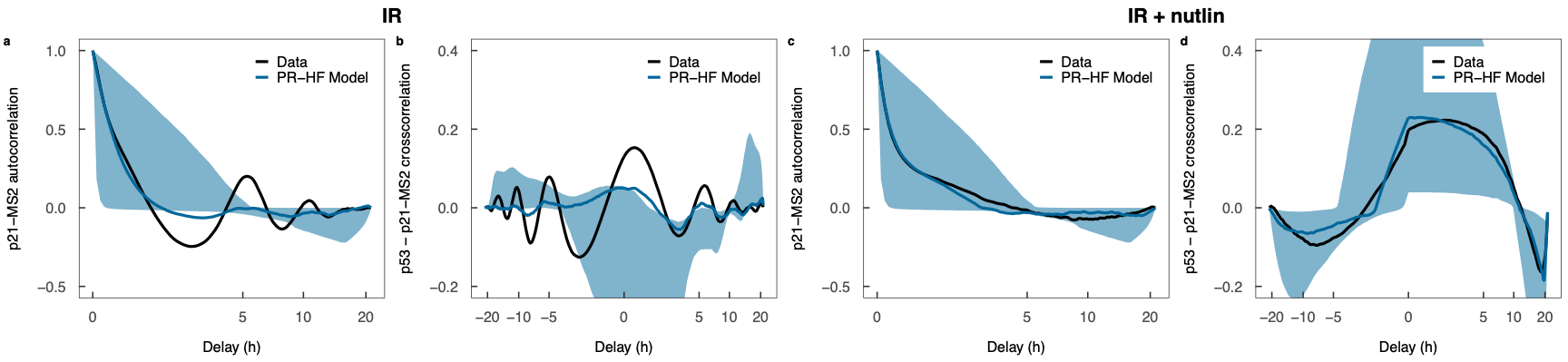

**Supplementary Figure 5: Comparison of experimental and simulated correlation functions when using the PR-Hill Function model. (a)** Mean p21-MS2 autocorrelation function for 10 Gy IR-treated cells for the PR-Hill Function model. **(b)** Mean p53 - p21-MS2 cross-correlation function for 10 Gy IR-treated cells for the PR-Hill Function model. **(c)** Mean p21-MS2 autocorrelation function for 10 Gy IR + 10 $\mu$M nutlin-3a-treated cells for the PR-Hill Function model. **(d)** Mean p53 - p21-MS2 cross-correlation function for 10 Gy IR + 10 $\mu$M nutlin-3a-treated cells for the PR-Hill Function model. In all panels, the colored lines represent the correlation functions from the simulations that are closest to the correlation functions from the data and the shaded areas represent the 95 percentile confidence intervals of the correlation functions from the simulations. Correlation functions are the means of 248 and 251 cells treated with 10 Gy IR and 10 Gy IR + 10 $\mu$M nutlin-3a, respectively.

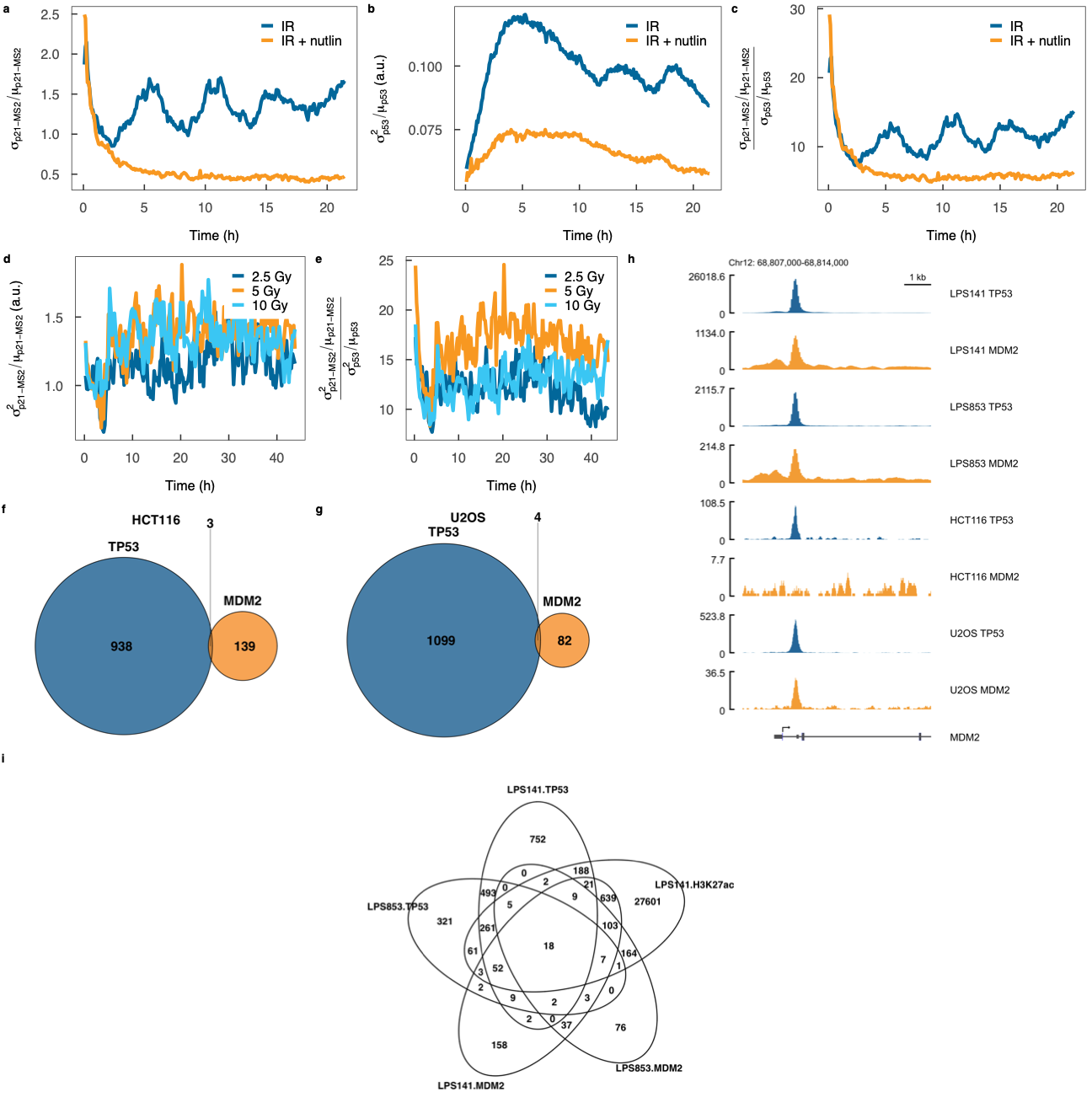

**Supplementary Figure 6: The incoherent feedforward loop increases p21 expression noise. (a)** Longitudinal measurements of p21-MS2 noise, as measured by the coefficient of variation, in cells treated with 10 Gy IR and 10 Gy IR + 10 $\mu$M nutlin-3a. **(b)** Longitudinal measurements of p53 protein noise, as measured by the Fano factor, in cells treated with 10 Gy IR and 10 Gy IR + 10 $\mu$M nutlin-3a. **(c)** Longitudinal measurements of p21-MS2 to p53 protein noise ratio, with noise measured by the coefficient of variation, in cells treated with 10 Gy IR and 10 Gy IR + 10 $\mu$M nutlin-3a. **(d)** Longitudinal measurements of p21-MS2 noise, as measured by the Fano factor, in cells treated with different doses of IR. **(e)** Longitudinal measurements of p21-MS2 to p53 protein noise ratio, with noise measured by the Fano factor, in cells treated with different doses of IR. **(f)** Venn diagram of TP53 and MDM2 ChIP-seq peaks in the HCT116 cell line. **(g)** Venn diagram of TP53 and MDM2 ChIP-seq peaks in the U2OS cell line. **(h)** TP53 and MDM2 ChIP-seq peaks at the *MDM2* gene in LPS141, LPS853, HCT116 and U2OS cells. **(i)** Venn diagram of TP53, MDM2 and H3K27ac ChIP-seq peaks in the LPS141 and LPS853 cell lines.

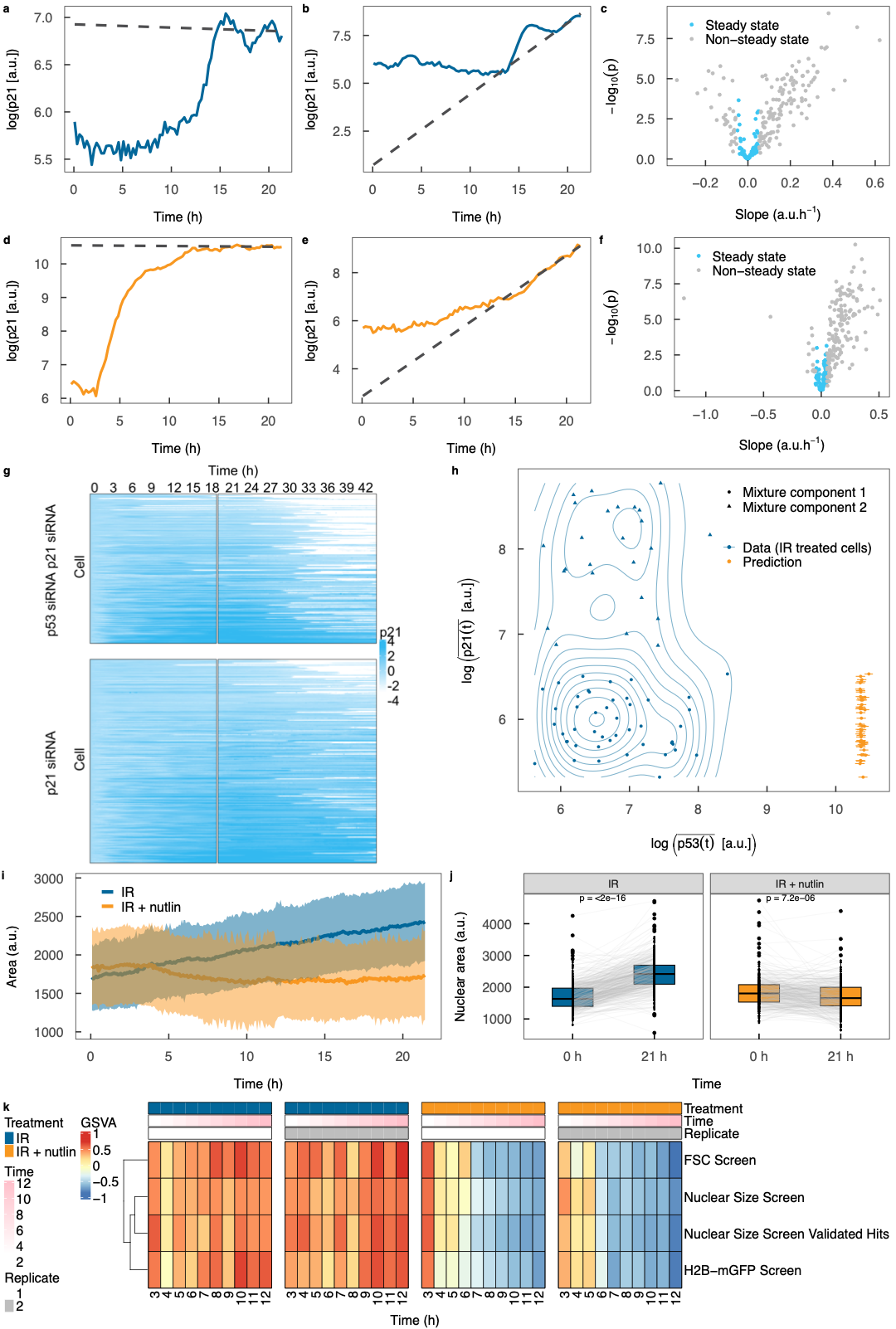

**Supplementary Figure 7: Abrogating MDM2-mediated transcriptional repression of p21 prevents the G1-S transition. (a)** p21 protein dynamics for a representative cell treated with IR reaching quasi-steady state. **(b)** p21 protein dynamics for a representative cell treated with IR not reaching quasi-steady state **(c)** Regression slope estimates (and p-values) for cells treated with IR used to define cells reaching steady state. **(d)** p21 protein dynamics for a representative cell treated with IR + nutlin-3a reaching quasi-steady state. **(e)** p21 protein dynamics for a representative cell treated with IR + nutlin-3a not reaching quasi-steady state. **(f)** Regression slope estimates (and p-values) for cells treated with IR + nutlin-3a used to define cells reaching steady state. In (a), (b), (d) and (e) the dotted line represents the regression fit to the final 2.5 h of the time course**. (g)** Single cell p21 dynamics following 10 Gy IR administration at the start of the experiment and p53 siRNA and p21 siRNA at 20 h (indicated by the column break) (n = 345). The color scale shows the log_2_ fold change in the fluorescence level and are truncated at the 5^th^ and 95^th^ percentiles. **(h)** Prediction of p21 protein levels from abrogating MDM2-mediated p53 degradation without altering the IFFL in IR-treated cells in S-phase. The orange points (and error bars) show the predictions (and their standard errors) and the blue points (and contours) show the data (and distribution) for IR-treated cells (same as in (Fig. 5A)). Abrogating MDM2-mediated p53 degradation without altering the IFFL is not predicted to increase p21 levels of S-phase cells to the levels of those in non-S-phase cells. **(i)** Nuclear area dynamics of 10 Gy IR- (n = 357) and 10 Gy IR + 10 $\mu$M nutlin-3a- treated (n = 313) MCF-7 cells extracted from timelapse microscopy. The lines indicate the mean and the shaded areas indicate 1 standard deviation from the mean. **(j)** Change in nuclear area in within single cells between the start and end of the experiment. p-values are from paired t-tests. **(k)** Gene set variation analysis of nuclear size gene sets^7^ applied to RNA-seq data measured hourly from 3 – 12 h following treatment of MCF-7 cells with 10 Gy IR or 10 Gy IR + nutlin-3a**.** GSVA – gene set variation analysis.

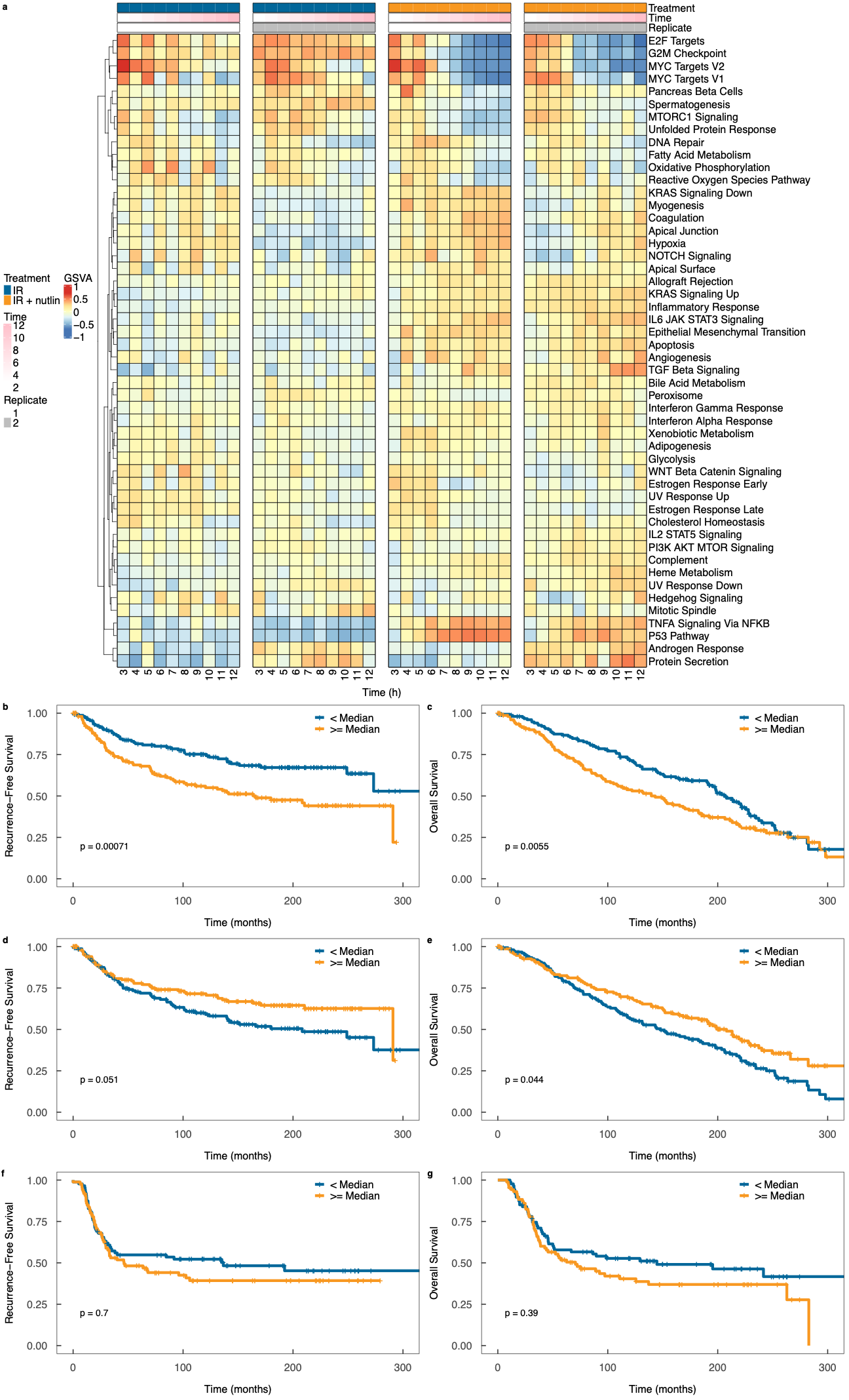

**Supplementary Figure 8: Abrogating the p53-MDM2 interaction steers cells into a persister state. (a)** Gene set variation analysis of Hallmark gene sets applied to RNA-seq data measured hourly from 3 – 12 h following treatment of MCF-7 cells with 10 Gy IR or 10 Gy IR + nutlin-3a. **(b)** Association between recurrence-free survival and GSVA score of genes downregulated in 10 Gy IR + nutlin-3a- versus 10 Gy IR-treated MCF-7 cells, in breast cancer patients not treated with adjuvant therapy from the METABRIC cohort (n = 311). **(c)** Association between overall survival and GSVA score of genes downregulated in 10 Gy IR + nutlin-3a- versus 10 Gy IR-treated MCF-7 cells, in breast cancer patients not treated with adjuvant therapy from the METABRIC cohort (n = 311). **(d)** Association between recurrence-free survival and GSVA score of genes upregulated in 10 Gy IR + nutlin-3a- versus 10 Gy IR-treated MCF-7 cells, in breast cancer patients not treated with adjuvant therapy from the METABRIC cohort (n = 311). **(e)** Association between overall survival and GSVA score of genes upregulated in 10 Gy IR + nutlin-3a- versus 10 Gy IR-treated MCF-7 cells, in breast cancer patients not treated with adjuvant therapy from the METABRIC cohort (n = 311). **(f)** Association between recurrence-free survival and GSVA score of genes upregulated in 10 Gy IR + nutlin-3a- versus 10 Gy IR-treated MCF-7 cells, in breast cancer patients treated with chemo-radiation therapy from the METABRIC cohort (n = 173). **(g)** Association between overall survival and GSVA score of genes upregulated in 10 Gy IR + nutlin-3a- versus 10 Gy IR-treated MCF-7 cells, in breast cancer patients treated with chemo-radiation therapy from the METABRIC cohort (n = 173). The p-values in (b) - (g) are from Cox proportional hazards regression models with the signature score, as a continuous variable, as the covariate.
